# Revealing the full biosphere structure and versatile metabolic functions in the deepest ocean sediment of the Challenger Deep

**DOI:** 10.1101/2021.06.05.447043

**Authors:** Ping Chen, Hui Zhou, Yanyan Huang, Zhe Xie, Mengjie Zhang, Yuli Wei, Jia Li, Yuewei Ma, Min Luo, Wenmian Ding, Junwei Cao, Tao Jiang, Peng Nan, Jiasong Fang, Xuan Li

**Affiliations:** Key Laboratory of Synthetic Biology, CAS Center for Excellence in Molecular Plant Sciences, Institute of Plant Physiology and Ecology, Chinese Academy of Sciences, Shanghai, China; Ministry of Education Key Laboratory for Biodiversity Science and Ecological Engineering, School of Life Sciences, Fudan University, Shanghai, China; Shanghai Engineering Research Center of Hadal Science and Technology, College of Marine Sciences, Shanghai Ocean University, Shanghai, China; University of Chinese Academy of Sciences, Beijing, China

**Author notes:** Corresponding author: Xuan Li, Jiasong Fang, Peng Nan. These authors contributed equally to this work (Ping Chen, Hui Zhou, Yanyan Huang, Zhe Xie, Mengjie Zhang).

**Keywords:** Mariana Trench, Challenger Deep, Hadal sediment, Metagenomics, Versatile metabolism, Metavirome, Large-scale cultivation

## Abstract

**Background:** The full biosphere structure and functional exploration of the microbial communities of the Challenger Deep of the Mariana Trench, the deepest known hadal zone on Earth, lag far behind that of other marine realms.

**Results:** We adopt a deep metagenomics approach to investigate the microbiome in the sediment of Challenger Deep, Mariana Trench. We construct 178 metagenome-assembled genomes (MAGs) representing 26 phyla, 16 of which are reported from hadal sediment for the first time. Based on the MAGs, we find the microbial community functions are marked by enrichment and prevalence of mixotrophy and facultative anaerobic metabolism. The microeukaryotic community is found to be dominated by six fungal groups that are characterized for the first time in hadal sediment to possess the assimilatory and dissimilatory nitrate/sulfate reduction, and hydrogen sulfide oxidation pathways. By metaviromic analysis, we reveal novel hadal Caudovirales clades, distinctive virus-host interactions, and specialized auxiliary metabolic genes for modulating hosts’ nitrogen/sulfur metabolism. The hadal microbiome is further investigated by large-scale cultivation that cataloged 1070 bacterial and 19 fungal isolates from the Challenger Deep sediment, many of which are found to be new species specialized in the hadal habitat.

**Conclusion:** Our hadal MAGs and isolates increase the diversity of the Challenger Deep sediment microbial genomes and isolates present in the public. The deep metagenomics approach fills the knowledge gaps in structure and diversity of the hadal microbiome, and provides novel insight into the ecology and metabolism of eukaryotic and viral components in the deepest biosphere on earth.

## Background

The hadal trench represents the deepest habitat for living organisms on the surface of the earth and accounts for a significant portion of the global benthic area [1]. In addition to elevated hydrostatic pressure (60-110 MPa), the trench environments are characterized as near-freezing temperatures, total darkness, poor nutrient availability, and isolation in topography [2]. Despite the harsh conditions, abundant microorganisms and metabolic activities were found to exist in both the hadal water columns and hadal sediments [3–5]. Hadal trenches were proposed to comprise specialized biodiversity related to their geographic isolation and unique environment characteristics [2]. The microbial abundance in hadal trenches was related to the availability of sedimentary organic matter, which reached the deep hadal environments by sinking via the funneling effect and by occasional landslides induced by deep ocean earthquakes [4–6].

A great deal of effort has been focused on the Mariana Trench system in the Western Pacific ocean where two tectonic plates, the Philippine Sea plate and the Pacific plate, collide [7, 8]. It contains the deepest habitat known on earth, the Challenger Deep, ∼11,000 meter below the ocean surface [9]. Characterization of the microbial species in the hadal habitat began in the 1950s using culture-based techniques [10–12]. Recent technological advances using 16S ribosomal RNA (rRNA) gene profiling expanded studies on ocean microbes by circumventing the limitation of culture dependency. Investigation on microbiome from abyssal water to bottom sediment in the Mariana Trench via the 16s rRNA gene profiling found that Proteobacteria, Bacteroidetes, Actinobacteria, Gemmatimonadetes, Thaumarchaeota, and Planctomycetes were dominant in the Mariana Trench habitats [4, 13, 14]. In a comparison of microbes between Mariana and Kermadec trench habitats, they comprised cosmopolitan taxa with different abundances, in addition to some autochthonous microbes associated with unique and rare OTUs [15, 16].

However, our understanding of the hadal microbial diversity and association of environmental factors has largely relied on the technology of 16s rRNA gene profiling, and the studies of limited number of cultured microbes. It was suggested that genetic and phenotypic details of trench microbial communities would ultimately require whole metagenome studies, but not just 16S rRNA gene analyses [16]. Despite the advance in sequencing technology, few high-throughput metagenomics studies on the microbial communities in the Challenger Deep sediment habitat have been conducted. Knowledge gaps remain about its biosphere structure, and details are sparse on the metabolic functions of different microbial components that drive the biogeochemical processes in the deepest habitat. Furthermore, currently little is known about the microeukaryotic and viral components of the hadal sediment biosphere, and essential questions about their community functions and ecological importance remain unanswered.

In the current study, a two-fold strategy was adopted to investigate the microbial community structure and metabolic functions in the Challenger Deep sediment, the deepest habitat on the earth. First, a deep metagenomics approach was designed and employed, by preparing extensive metagenomic libraries for Illumina sequencing, to achieve unprecedented coverage depth on its metagenome. This strategy is necessary to capture microbes of low abundances and unravel the full community structure of the hadal biosphere, underpinning its eukaryotic and viral components. It enabled us to reconstruct the largest dataset of metagenomic assembled genomes (MAGs) and to identify the versatile metabolic functions of the hadal microbiome in great detail. Second, large-scale cultivation was conducted to isolate microorganisms from the Challenger Deep sediment biosphere, using twenty-four different types of media in combination with different culture conditions. We obtained more than 2000 microbial isolates and cataloged 1070 bacteria and 19 fungi by 16S rRNA gene or nuclear ribosomal internal transcribed spacer (ITS) tag sequencing. These microbial isolates became valuable resources for further study of adaptive mechanisms in extreme habitat.

## Results and discussion

### Hadal sediment geochemistry and deep metagenomic sequencing

Sediment samples were collected using two deep-sea hadal landers from the seafloor (depth of 10840 meters) at the Challenger Deep (142°21.7806’E, 11°25.8493’N). The sediment samples were dissected into three depth segments, i.e. the surficial segment MT-1 (sediment depth 0-5 cm), the mid-segment MT-2 (5-10 cm), and the deep segment MT-3 (10-14 cm). Geochemical measurements on the samples were taken either onboard the ship ZhangJian or inland laboratories using preserved samples. The contents of total organic carbon (TOC) and total nitrogen (TN) in the sediment were measured between 0.49 and 0.55 (wt%) and between 0.05 and 0.06 (wt%), respectively (Additional file 1: Table S1). They are within the previously reported values for the sediment of the Challenger Deep [13]. The values of δ^13^C (-21.41 to -21.53 ‰) and δ^15^N (5.42 to 6.69 ‰) were within the ranges of the commonly observed values for marine organic matter [17]. This agrees with the results of recent studies that marine algae were the dominant source of sedimentary organic matter in the southern Mariana Trench [18, 19]. For nutrient ions, while the porewater SO_4_^2-^ concentrations were constant throughout the three segments, those of NH_4_^+^, PO_4_^3-^, and NO_2_^-^ were trending upward with increasing sediment depth (Additional file 1: Table S2). The NO_3_^-^ concentrations, on the other hand, decreased with depth. The dissolved major and trace elements remained relatively homogenous with little variations among the three segments (Additional file 1: Table S3 and S4).

To explore the full microbiome structure in the Challenger Deep sediment, we performed deep metagenomic sequencing on the sediment samples which was designed with enhanced sensitivity, to capture the genetic contents of microbes with low abundances. Note that we took steps to minimize the impact of sample temperature increase by swiftly processing and freezing sediment samples, preserving the big-picture characteristics of the microbiome for metagenomic analysis. Three or more independent Illumina libraries were generated for each segment, and on average 22.6 Gb sequence data were obtained from each library, generating a total of 248.65 Gb raw sequence data (Table S5). Metagenomic co-assembly was carried out using all the clean data, which resulted in a metagenome of 6.65 Gb in total length with ∼6.21 million contigs and an N50 of ∼1.14 Kb.

To reveal the compositions of the hadal sediment communities, taxonomic profiling analysis was performed on the metagenomics sequences using the kaiju program and the NCBI nr library [20, 21]. The relative sequence abundance for bacteria and archaea in the sediment accounted for 82.07% and 6.23%, respectively, whereas that for microeukaryotes and viruses was 0.69% and 0.12%, neither of which has been reported for the Challenger Deep sediment before. Different from previous studies based on rRNA gene PCR-tag sequencing, our approach estimated the sequence abundance values via the same workflow, generating scores directly comparable within the community scope.

At the phylum level, the most abundant components were Proteobacteria, Chloroflexi, Actinobacteria, Thaumarchaeotathat, Planctomycetes, Firmicutes, Bacteroidetes, Gemmatimonadetes, Acidobacteria, and Gemmatimonadetes, belonging to either bacteria or archaea (Fig. 1A). Our results agree with previous studies relying on 16S rRNA PCR-amplicon sequencing, in which Alphaproteobacteria, Chloroflexi, and Gemmatimonadetes were the most abundant bacteria, and Thaumarchaeota the most abundant archaea found in the Challenger Deep sediment habitat [16, 22]. However, the relative abundance of Thaumarchaeota varied largely between the different studies of the Mariana Trench habitats, ranging from 0.67% to 67% [23] and from 0.5% to 40% [22]. In a non-Mariana system, the Yap Trench, while Thaumarchaeota was similarly enriched, the abundance of Chloroflexi, Actinobacteria, Planctomycetes, and Gemmatimonadetes in its sediment microbiome was significantly lower than that of Mariana Trench sediment habitats [24]. We found that the overall relative abundance of archaea decreased with sediment depth, similar to findings in a previous study [13]. However, the relative abundance of both the microeukaryotes and marine viruses increased with sediment depth. The most abundant eukaryotes were Opisthokonta (Fungi), Alveolata, Stramenopiles, and Rhodophyta. Note the top eukaryotic phyla, Ascomycota and Basidiomycota, ranked the 16th and 17th overall in sequence abundance (Fig. 1A). We found MT-1 and MT-2 had consistent microbial compositions compared to those of MT-3, suggesting a geochemical boundary separated them at a 10-cm depth that was previously shown to limit oxygen access in hadal sediment [25]. Albeit in a relatively low abundance, the eukaryotic and viral components were unraveled for the first time in the Challenger Deep sediment, and as integral parts of the hadal biosphere, their ecological significance was further addressed below.

**Fig. 1.**
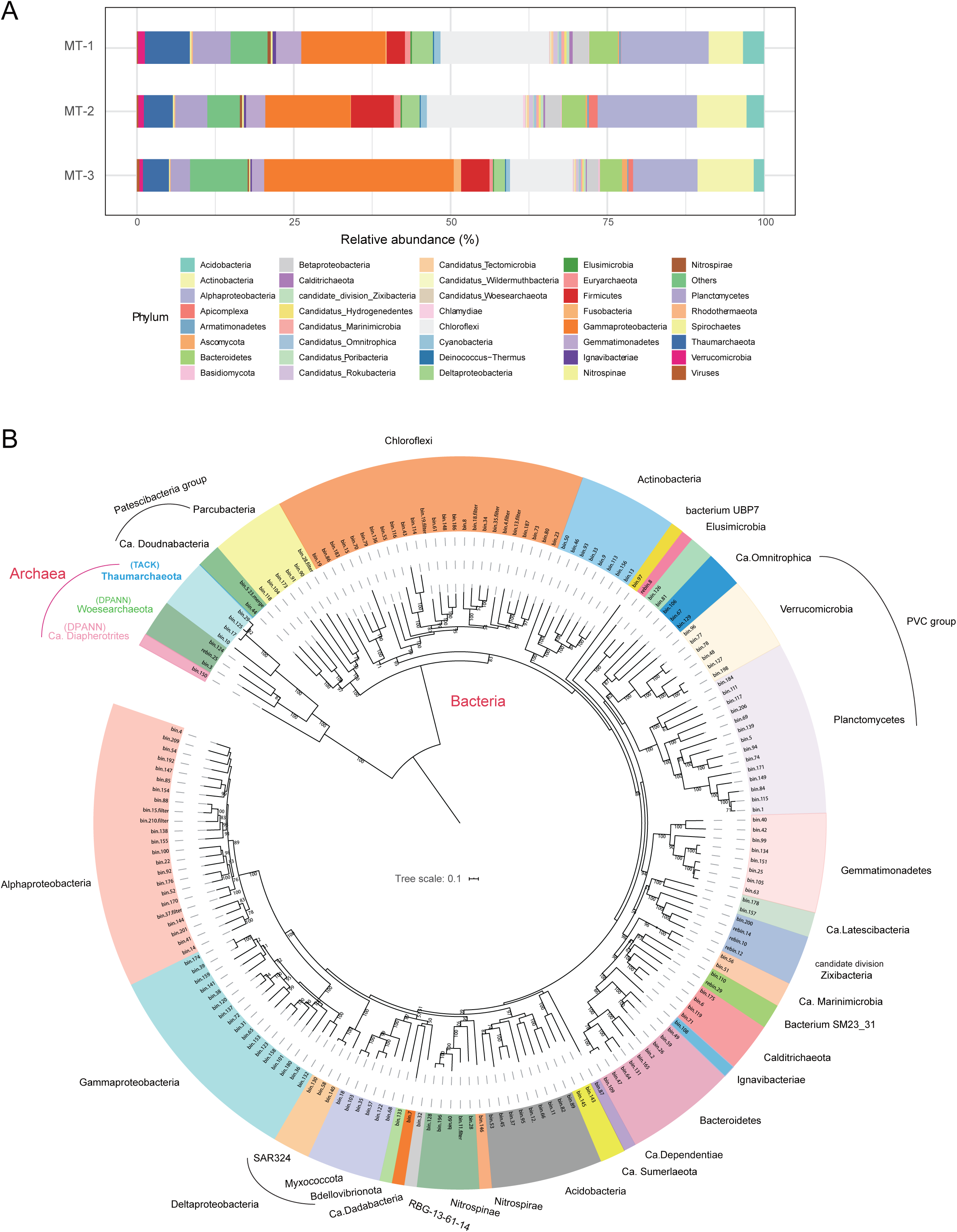
Composition and phylogenetic diversity of the microbial community in the Challenger Deep sediment habitat. (A) The relative sequence abundance of dominant microbial groups (top 30). (B) Phylogenetic analysis of the 178 MAGs based on 43 conserved single-copy, protein-coding marker genes using the maximum likelihood algorithm. Bootstrap values based on 1000 replications are shown for each branch. The scale bar represents 0.1 amino acid substitution per position.

### Phylogenetic diversity of the hadal sediment microbiome

Metagenomic binning on the assembled metagenome and filtering resulted in 178 quality draft genomes, i.e. MAGs (metagenome-assembled genomes) (Additional file 1: Table S6). These MAGs were at or above the medium-quality standards for draft genome, which required >50% completeness and <10% contamination (*Methods*) [26]. Furthermore, among them, 101 draft genomes were >70% completeness, and 22 were >90% completeness. The MAGs represented the most diverse metagenome reconstructed from the hadal sediment biosphere, comprising members from 26 phyla or candidate phyla. Sixteen of them had MAGs found from the Changer Deep sediment for the first time, i.e. Gemmatimonadetes (8), Verrucomicrobia (8), Nitrospinae (5), Elusimicrobia (1), Ignavibacteriae (1), Calditrichaeota (4), candidate phyla including Zixibacteria (4), Ca. Hydrogenedentes (2), Ca. Dependentiae (1), Ca. Diapherotrites (1), Ca. Doudnabacteria (2), Ca. Latescibacteria (2), Ca. Omnitrophica (3), Ca. Marinimicrobia (2), and unclassified bacteria including Bacterium SM23_31 (2), Ca. Sumeriaeota (2) and a newly defined phylum UBP7_A (1) [27]. In comparison, previous studies reported eleven and thirty MAGs that were co-assembled by combing water and sediment microbial sequences from the Mariana Trench and were affiliated with three and twelve phyla, respectively [28, 29]. Besides, using the single-cell sequencing approach, twelve single-cell amplified genomes (SAGs) were generated for Parcubacteria from the Mariana Trench sediment [30]. Note that these earlier works had fewer MAGs despite co-assembly by mixing sample data from multiple sources. Thus, the deep metagenomics approach has significantly enhanced the coverage and sensitivity of the hadal microbiome.

To illustrate the taxonomic diversity of the hadal sediment microbes, a phylogenetic tree for the 178 MAGs was constructed (Fig. 1B), using 43 conserved single-copy, protein-coding marker genes [31, 32]. A substantial number of uncultured microbial lineages were uncovered and classified with the phylogenetic analysis. The main bacterial groups included Alphaproteobacteria (22 MAGs), Gammaproteobacteria (19), Chloroflexi (19), Planctomycetota (14), Bacteroidota (10), Actinobacteriota (10), Gemmatimonadetes (8), Verrucomicrobia (6), Nitrospinae (4), as well as Elusimicrobia (1), Ignavibacteriae (1), Nitrospirae (1) and Calditrichaeota (4). Notably, bin.97 formed a monophyletic clade in the phylogenetic tree, and was distant from the PVC groups that include Planctomycetes (Fig. 1B). It was closely related to Planctomycetales bacterium 4484_113 based on analysis using GTDB-Tk [33], for which the Average Nucleotide Identity (ANI) to bacterium 4484_113 stood at 63.86%. Bacterium 4484_113 was originally defined as unclassified Planctomycetes, but a recent study separated it from Planctomycetes to form a new phylum UBP7_A [27]. Thus, our study uncovered a likely second member of UBP7_A from the Challenger Deep sediment habitat.

The main archaea groups included Ca. Woesearchaeota (3 MAGs), Thaumarchaeota (4), and Ca. Diapherotrites (1) (Fig. 1B). Note that Ca. Diapherotrites, represented by bin.150, was found for the first time in the Challenger Deep sediment. Intriguingly, bin.150 was situated as an outgroup to both Thaumarchaeota (TACK group) and Woesearchaeota, and had a closer relationship with bacteria in the phylogenetic tree (Fig. 1B). Based on analysis using GTDB-Tk, bin.150 was close to the unclassified Diapherotrites archaeon UBA493 that is affiliated to DPANN group. Its ANI to UBA493, the closest relative, was 72.19%. We placed bin.150 with additional members of Ca. Diapherotrites for phylogenetic analysis, and found bin.150 formed a long-branch that represents a specialized Diapherotrites clade in the Challenger Deep sediment (Additional file 2: Fig. S1). A previous study reported that as a member of Ca. Diapherotrites, Ca. Iainarchaeum acquired anabolic genes from bacteria via horizontal gene transfer [34]. The same reason may explain why bin.150 had a close relationship with bacteria in our phylogenetic results. So by reconstructing the largest metagenome of the hadal sediment biosphere, we recovered representative genomes of all major prokaryotic lineages previously identified by 16S rRNA gene amplicon-based surveys for the Challenger Deep sediment habitat [35], providing valuable references for us to further look into details of hadal microbiome regarding genetic diversity, metabolic functions, and symbiotic relationship. While our analyses were based on constructed MAGs, we acknowledged the missing pathway components that are likely attributed in part by the incomplete assembly, and the many assembled sequences of unknown molecular functions that are knowledge gaps to be bridged with new means [36].

### Versatile metabolic function of the hadal sediment microbiome

#### Heterotrophy vs. Autotrophy

To investigate the metabolic potential of each component, the 178 MAGs reconstructed from the Challenger Deep sediment microbiome were assigned with metabolic functions based on KEGG annotation. To determine microbes’ lifestyle, the MAGs were first investigated for heterotrophic potential and were found to contain genes involved in various pathways for degradation of carbohydrates (all MAGs), hydrocarbons (34 MAGs affiliated with 9 phyla), and aromatic compounds (146 MAGs affiliated with 24 phyla) (Fig. 2 and Additional file 1: Table S7). It is likely that sinking particulates from the upper ocean or terrestrial inputs, partly due to the funneling effect and earthquake-inducing landslides, were the source of the organic matter in the deepest habitat [37].

**Fig. 2.**
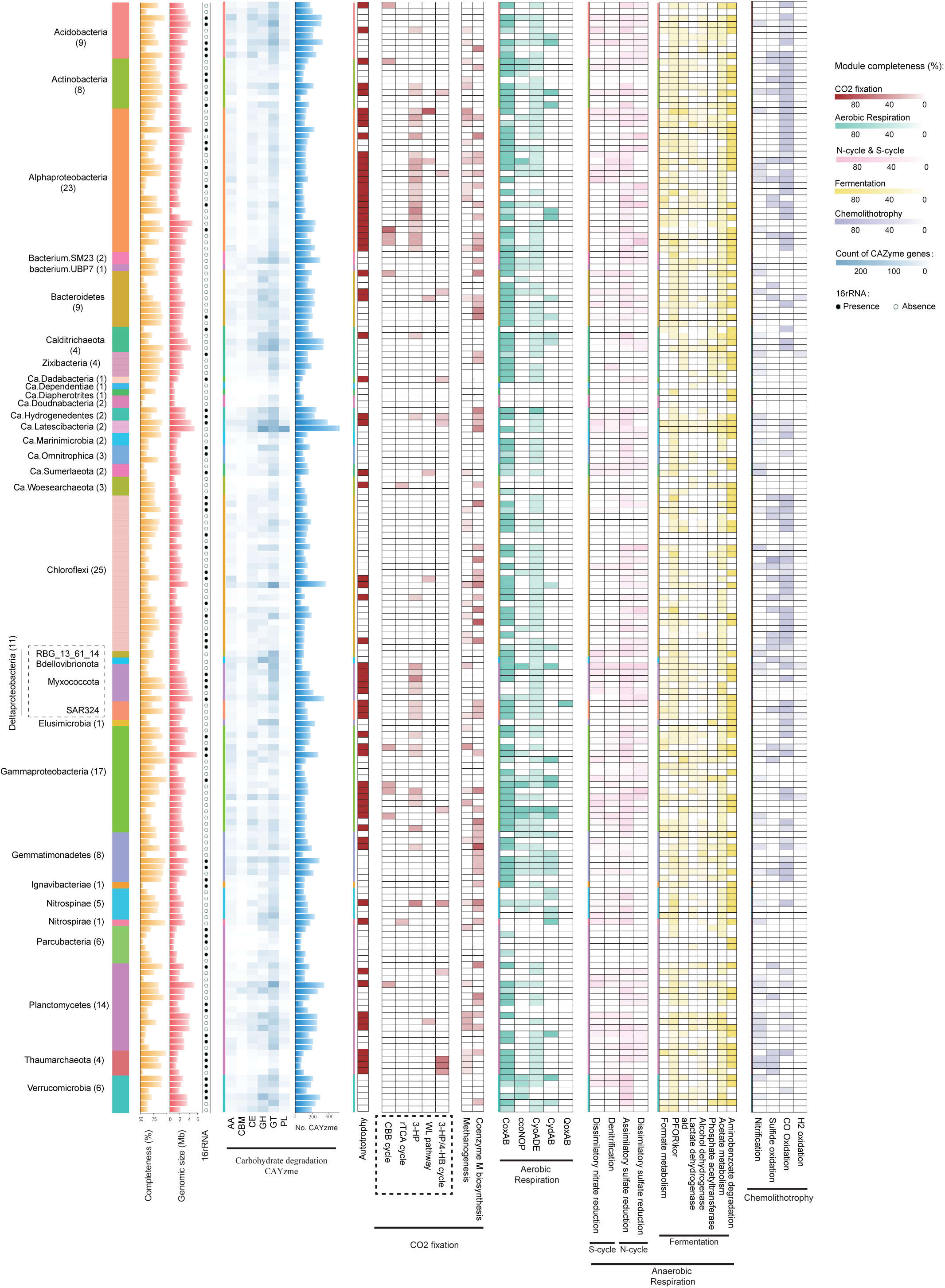
Heat-map presentation of genomic features and metabolic potential for the 178 MAGs (with the taxonomic assignment) reconstructed for the Challenger Deep sediment microbiome. Key genes involved in Carbohydrate degradation (CAYzme), CO_2_ fixation, Aerobic Respiration, Anaerobic Respiration, and Chemolithotrophy are illustrated (refer to Additional file 1: Table S7 for details). Abbreviations: GH, glycosidases or glycosyl hydrolases; PL, polysaccharide lyases; CE, carbohydrate esterases; GT, glycosyltransferases; AA, auxiliary activities; CBM, carbohydrate-binding modules; WL, Wood-Ljungdahl pathway; CBB, Calvin-Benson-Bessham cycle; rTCA, reverse tricarboxylic acid cycle; 3-HP: 3-hydroxypropionate bi-cycle; 3-HP/4-HB cycle, 3-hydroxypropionate/4-hydroxybuty rate cycle.

On the other hand, we found 69 out of 178 MAGs (39%; 16 phyla) contained pathways for inorganic carbon fixation. Six different CO_2_ fixation mechanisms were found to involve various microbes (Fig. 2). Among them, 3-hydroxypropionate bi-cycle (3-HP) was the most prevalent in 43 MAGs, compared to the Calvin-Benson-Bassham Cycle (CBB or Calvin cycle) (11 MAGs), Wood-Ljungdahl pathway (WL pathway) (6 MAGs), the reverse TCA cycle (rTCA) (2 MAGs), the 3 hydroxypropionate-4 hydroxybutyrate cycle (3-HP/4-HB cycle) (6 MAGs), and methanogenesis (16 MAGs). Previously, 3-HP was found in Chloroflexaceae, Alpha- and Gammaproteobacteria, and SAR202 [38, 39]. In our study, the 43 MAGs detected to have the 3-HP pathway were taxonomically assigned to eight phyla (31%) (Fig. 2 and Additional file 1: Table S7). They include the unclassified Chloroflexi bacterium RBG_16_64_32, Alpha-, Delta-, and Gamma-proteobacteria, Nitrospinae, Acidobacteria, Actinobacteria, Calditrichaeota, Ca. Hydrogenedentes, and Gemmatimonadetes.

For other CO_2_ fixation pathways, the key enzymes in the Calvin Cycle, ribulose-bisphosphate carboxylase (*rbcS*), and phosphoribulokinase (*prkB*) were detected in five phyla (Fig. 2 and Additional file 1: Table S7). The rTCA cycle biomarker *Acl* was present in one MAG for Nitrospira and one MAG for Ca. Woesearchaeota. Further, rTCA cycle was the only CO_2_ fixation pathway employed by Nitrospira and Ca. Woesearchaeota. While nitrite-oxidizing Nitrospira was previously known to use rTCA cycle for CO_2_ fixation [40], this is the first reported case that Ca. Woesearchaeota may be capable of rTCA reaction. 3-HP/4-HB cycle was the most energy-efficient aerobic autotrophic pathway for CO_2_ fixation[41]. The biomarker enoyl-CoA hydratase/3-hydroxyacyl-CoA dehydrogenase involved in 3-HP/4-HB or DC/4-HB cycle, was found in three bacterial groups, Nitrospinae, Acidobacteria and Planctomycetes, and one archaea, Thaumarchaeota. The WL pathway biomarkers, i.e. acetyl-CoA decarbonylase/synthase complex subunit delta (*cdhD*), acetyl-CoA synthase (*acsB*), anaerobic carbon-monoxide dehydrogenase (*cooS* and *cooF*), and 5-methyltetrahydrofolate corrinoid/iron-sulfur protein methyltransferase (*acsE*) were found in Bacteroidetes, Alphaproteobacteria, Ca. Sumerlaeota, Chloroflexi, Gammaproteobacteria, and Planctomycetes. Moreover, we found 16 MAGs associated with 7 phyla, i.e. Proteobacteria, Bacteroidetes, Chloroflexi, Gemmatimonadetes, Nitrospinae, Planctomycetes, and Thaumarchaeota, contained the Coenzyme M biosynthesis and methanogenesis genes (Fig. 2 and Additional file 1: Table S7), indicating the hadal microorganisms had the potential to produce methane from fixing CO_2_ in the hadal zone.

Overall, among the 178 MAGs reconstructed for the hadal sediment microbiome, 69 MAGs (∼39%) were affiliated with 16 phyla and mixotrophic based on their capacity of inorganic carbon fixation (Fig. 2 and Additional file 1: Table S7). The enrichment and significant taxonomic expansion of mixotrophic microbes we revealed for the deepest hadal microbiome, may indicate an adaptive niche that mixotrophy confers to microbes living in an oligotrophic habitat like the Challenger Deep sediment.

#### Aerobic vs Anaerobic Respiration

We investigated the hadal microbiome for its potential to carry out aerobic or anaerobic respiration. 155 MAGs (∼87%) were found to contain aerobic respiratory genes, such as Cytochrome c oxidases (Cox/Cyd/Qox/cco/Cyo). These MAGs are associated with 24 phyla, such as Acidobacteria, Bacteroidetes, Gemmatimonadetes, and Proteobacteria (Fig. 2 and Additional file 1: Table S7). Therefore, a large majority of the microbes in the hadal habitat can potentially use oxygen as an electron acceptor for energy generation.

On the other hand, anaerobic respiration pathways were also found to be widely distributed, for which nitrate/nitrite or sulfate was the alternative electron acceptors. Dissimilatory nitrate reduction to ammonia pathway (DNRA), denitrification pathway, and sulfate reduction pathways were searched and found in a total of 145 MAGs (81%) (Fig. 2 and Additional file 1: Table S7). These MAGs covered 21 of the 26 phyla we identified in the hadal microbiome. Specifically, the genes for DNRA (*nirB* and *nirD* as markers), denitrification pathway (*nirK* and *norC*), assimilatory sulfate reduction pathway (*sat, aprAB* and *dsrAB*), and dissimilatory sulfate reduction pathway (*aprA-* and *aprB*) were present in 64 MAGs (15 phyla), 77 MAGs (16 phyla), 135 MAGs (20 phyla), and 67 MAGs (15 phyla), respectively (Fig. 2 and Additional file 1: Table S7). Previous studies based on 16S rRNA gene sequencing analysis suggested that dissimilatory nitrate reduction and denitrification were the dominant reactions, whereas microbial sulfate reduction was negligible in the Challenger Deep sediment [13]. However, our analysis shows that the broad distribution of both assimilatory sulfate reduction and dissimilatory sulfate reduction points to the equal importance of sulfate reduction pathways for anaerobic respiration, if not otherwise more important. In addition, for microbes capable of anaerobic respiration, many were found to have the potential for different types of fermentation for degradation of organic matters (Additional file 3: Analysis of fermentation; Additional file 1: Table S7).

The enrichment of anaerobic or anaerobic respiration metabolism in microbial community was apparently driven by the local habitat conditions. Examples of the deep petroleum seeps sediments and Guaymas Basin hydrothermal sediments illustrated MAGs enriched for anaerobic respiration in the presence of hydrocarbons i.e. alkanes and aromatic compounds, or depleted oxygen levels in sediments [31, 42]. We further investigated the MAGs for the potential to carry out both aerobic and anaerobic respiration, and found 136 MAGs out of 178 (∼76%), affiliated with 20 phyla (Fig. 2 and Additional file 1: Table S7), were capable of facultative anaerobic metabolism. Compared to the microbial communities in deep petroleum seeps or Guaymas Basin hydrothermal habitat, the high proportion of facultative anaerobes in the Challenger Deep sediment, reflects the microbial metabolic versatility and the ability to adapt by the hadal microbes to endemic habitats or environmental disturbance.

#### Chemolithotrophy

Microbes are known to acquire their energy for growth and CO2 fixation from the oxidation of inorganic compounds, such as hydrogen (H_2_), hydrogen sulfide (H_2_S), ammonia (NH_3_), carbon monoxide (CO), and metals[43]. The nitrification gene markers were found in 53 MAGs (30%, 17 phyla) of the Challenger Deep sediment microbiome (Fig. 2 and Additional file 1: Table S7). The key enzyme in nitrite oxidation, nitrite oxidoreductase/nitrate reductase was present in 35 MAGs (20%; 13 phyla), suggesting that nitrification is an important approach for energy production for a substantial proportion of microbes in the hadal sediment biosphere. Other genes involved in nitrification pathways, *PmoA-amoA*, *PmoB-amoB* and *PmoC-amoC*, were detected in 4 MAGs associated with 3 phyla (Additional file 1: Table S7). Hydroxylamine dehydrogenase gene (*hao*) was detected in 21 MAGs (12%; 12 phyla). Interestingly, the gene, *hao*, was not found in bathypelagic microbes capable of nitrification[44], reflecting the difference in microbial mediators between the bathypelagic and hadal zones in similar biogeochemical process.

The genes for CO-oxidation, sulfide-oxidation, H_2_-oxidation, and iron-oxidation were also detected in the Challenger Deep sediment microbiome. Carbon monoxide dehydrogenase (CODH; *cox* gene) was found in 92 MAGs (52%; 15 phyla) (Fig. 2 and Additional file 1: Table S7), indicating its broad distribution, and the importance of CO-oxidation as an energy source in the hadal habitat. Sulfide oxidation genes such as *sqr* and *fccB*, were found in 33 MAGs (19%; 8 phyla), i.e. Acidobacteria, Actinobacteria, Bacteroidetes, Chloroflexi, Gemmatimonadetes, Planctomycetes, Thaumarchaeota, and Proteobacteria. For iron oxidation, we detected *cyc2* gene (encoding Cytochrome c) in one MAG associated with Gemmatimonadetes, suggesting that Fe^2+^ can serve as an alternative electron donor for Gemmatimonadetes.

### Microeukaryotic community and predominant fungal groups

Microbial eukaryotes in hadal sediment were the least studied and remained largely unknown. Our analysis found that Opisthokonta (Fungi) were the predominant eukaryotes in the Challenger Deep sediment, accounting for 87.42% of the total eukaryotic sequences. Ascomycota (41.73%), Basidiomycota (26.82%), and Mucoromycota (12.64%) were the top phyla, followed by Chlorophyta (6.61%), Rhodophyta (2.44%), Bacillariophyta (2.31%), Chytridiomycota (2.21%), Zoopagomycota (1.91%), Apicomplexa (0.96%), etc (Fig. 3A). Our results revealed the presence of Stramenopiles e.g. Blastocladiomycota, and Alveolata e.g. Apicomplexa, two members of the super-group SAR (i.e. Stramenopiles, Alveolata, and Rhizaria), in the hadal sediment, albeit at low abundances. Rhizaria was, however, notably absent. Previous studies showed that the abundance of Alveolata and Stramenopiles decreased with depth in the water column of the Mariana Trench, whereas Opisthokonta had an inverse trend [45]. Our study expanded the findings into the trench sediment, illustrating the co-existence pattern of Alveolata, Stramenopiles, and Opisthokonta.

**Fig. 3.**
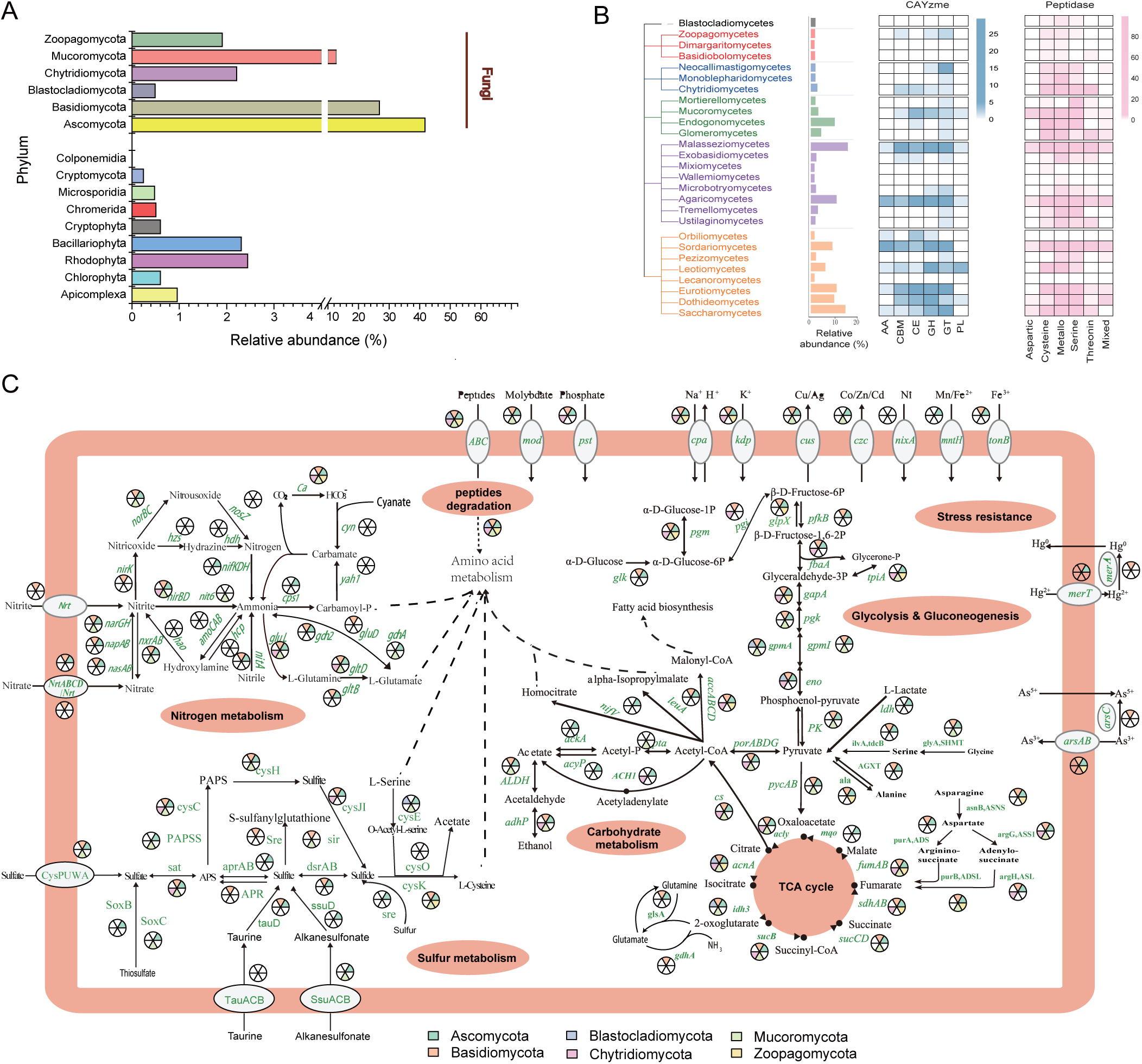
Composition of microeukaryotic community, and metabolic functions of the dominant fungal groups in the Challenger Deep sediment habitat. (A) The relative sequence abundance of different eukaryotic groups within total eukaryotes. (B) Phylogenetic relationship of major fungal groups identified in the hadal habitat, and the profiles of a carbohydrate-active enzyme family (CAZymes) and peptidase family genes. The numbers of genes detected are denoted by shade intensity. Abbreviations: GH, glycosidases or glycosyl hydrolases; PL, polysaccharide lyases; CE, carbohydrate esterases; GT, glycosyltransferases; AA, auxiliary activities; CBM, carbohydrate-binding modules. (C) Metabolic potentials of carbon, nitrogen, sulfur, and iron metabolism shown for the six dominant fungal groups. The presence of genes within the metabolic pathways is denoted for each group by the fan areas with color for the corresponding phylum. Gene symbols and metabolites are labeled with the KEGG designation (refer to Additional file 1: Table S8 for details).

As the dominant eukaryotic group, fungi in the Challenger Deep sediment biosphere comprised six phyla, i.e. Zoopagomycota, Mucoromycota, Ascomycota, Basidiomycota, Chytridiomycota, and Blastocladiomycota, which can be further classified into twenty-seven classes (Fig. 3B). Fungi were known to be distributed globally in deep-sea water, including that in the Mariana Trench, as Basidiomycota and Ascomycota were the primary components of the deep-sea fungal community [46, 47]. Basidiomycota and Ustilaginomycetes were previously detected in deep-sea sediment using a cloning library method [48]. The capability of living in anoxia conditions may be crucial for fungi to adapt to the hadal sediment environments, as Ascomycetes, Basidiomycetes, and Chytridiomycetes were reportedly capable of fermentation and anaerobic growth in the deep ocean [49].

### Metabolic functions of the fungal community in the hadal sediment biosphere

The details of gene contents and metabolic functions of the fungal community in both the water column and deep-sea sediment remained largely unknown, as early studies were based on 18S rRNA gene/ ITS-amplicon sequencing [45, 47]. In the current work, we explored the metabolic potential of the fungal groups using the assembled metagenome.

#### Carbon metabolism

We investigated the potential of the hadal fungi to metabolize carbohydrates and peptides in the hadal sediment by looking into genes for carbohydrate-active enzymes (CAZYmes) and peptidases. In total, we detected 262 CAZYmes (88 family) and 703 peptidases (77 family) (Fig. 3B and Additional file 1: Table S8). Members of CAZyme included 79 GHs (glycosidases or glycosyl hydrolases), 8 PLs (polysaccharide lyases), 51 CEs (carbohydrate esterases), 83 GTs (glycosyltransferases), 14 AAs (auxiliary activities; associated with polysaccharide and lignin degradation), and 27 CBM (carbohydrate-binding modules). The major sources of CAZymes were Eurotiomycetes (number of genes n = 20), Agaricomycetes (n = 16), Sordariomycetes (n = 38), and Malasseziomycetes (n = 30). Particularly, key CAZyme genes encoding xylanase (GH10) were found in Agaricomycetes, and those encoding cellulase (GH5) were found in Agaricomycetes, Chytridiomycetes and Microbotryomycetes (Additional file 1: Table S8).

Peptidase genes were abundant in Endogonomycetes (n = 124), Agaricomycetes (n = 84), Sordariomycetes (n = 70), and Malasseziomycetes (n = 67) (Fig. 3B and Additional file 1: Table S8). Six peptidase groups, i.e. Aspartic (A), Cysteine (C), Metallo (M), Mixed (P), Serine (S) and Threonine (T) peptidases, were found in the fungal genomes. Among them, the most abundant was Metallo peptidase, consisting of 230 members that belong to 30 families, followed by Cysteine (n = 225; 21 families), Serine (n = 168; 19 families), Threonine (n = 43; 3 families), and Aspartic peptidases (n = 14; 3 families) (Fig. 3B and Additional file 1: Table S8). So, the hadal fungi have broad potentials in organic carbon cycling, capable of degrading a variety of carbohydrate and peptide substrates.

#### Nitrogen metabolism

The hadal sediment fungi were found to possess the complete pathways for dissimilatory nitrate reduction and assimilatory nitrate reduction, and the partial pathway for nitrogen denitrification, but lacked the ability for nitrification or anammox (Fig. 3C). The key enzymes for nitrate reduction, namely *NarGHI/NapAB* and *NirBD* for dissimilatory pathway, and *NR/NasAB* and *NIT-6* for assimilatory pathway, were found in several fungal groups, indicating dissimilatory nitrate reduction is an important function for the hadal fungi. Ammonia in hadal water could be generated from decomposition of nitrogenous organic matter [4]. However, it can be limited in hadal sediment. Therefore, the nitrate reduction by the sediment fungi can be an important source of ammonia for hadal sediment microbes.

The partial pathway for nitrogen denitrification was also found in the hadal sediment fungi, missing the *NosZ* gene that coverts nitrous oxide to nitrogen. Downstream of the *NarGHI/NapAB* genes (shared with the dissimilatory nitrate reduction pathway), the *NirK* and *NorBC* genes completed the pathway to covert nitrite to nitric oxide, and nitric oxide to nitrous oxide, which may be released into the hadal sediment environment (Fig. 3C). However, unlike the bacterial community, the fungal community appeared lacking the capability of nitrification, as some key enzymes were generally missing in nitrification and anammox pathways (Fig. 3C). Notably, denitrification pathway enzymes were found in Fusarium, Penicillium, and Aspergillus from the Challenger Deep sediment (Additional file 1: Table S8). Some Fusarium species in the deep-sea oxygen-limiting environments were reportedly capable of denitrification [50]. In other low-oxygen habitats, such as anaerobic marine sediment, salt-tolerant Penicillium and Aspergillus species were also identified to carry out the denitrification process [50]. These data illustrate the potential of the hadal fungi in nitrogen metabolism, possibly having a significant role in the ecological process in the hadal environment.

#### Sulfur metabolism

Sulfate reduction is one of the main anaerobic respiratory pathways that many microbes living in anaerobic conditions depend on. The hadal fungi in the Challenger Deep sediment were found to possess the complete pathways for assimilatory sulfate reduction, dissimilatory sulfate reduction, and sulfide oxidation (Fig. 3C). They were also found to contain partial pathway for the SOX system, but were incomplete for oxidation of thiosulfate. The data suggest that hadal sediment fungi - like hadal bacteria, could take part in sedimentary sulfur cycle, which has not been reported for fungal communities in deep sea sediment.

The possession of complete sulfate reduction enzymes indicated the hadal fungi had the potential of using sulfate reduction for energy production when living under anaerobic conditions (Fig. 3C). A high copy number of genes for sulfate reduction enzymes, like *Sat, CysC, CysH*, and *CysJ*, were detected in the hadal sediment fungi (Additional file 1: Table S8). The sulfate adenylyltransferase gene (*sat*) for catalyzing the bidirectional reactions between APS and sulfate was widely distributed in members of different phyla, e.g. Ascomycota (4 classes), Basidiomycota (3 classes), Chytridiomycota (2 classes), and Mucoromycota (4 classes).

In the sulfide oxidation pathway, oxidation of sulfide to sulfite was catalyzed by *DsrA/B*, whose genes were detected in several classes of Basidiomycota and Ascomycota, like Malasseziomycetes and Sordariomycetes. Subsequently, sulfite was oxidized to APS by adenylylsulfate reductases. Dothideomycetes (belonging to Ascomycota) contained *Apr-A* gene for sulfite oxidization to APS. Agaricomycetes (belonging to Basidiomycota) had the adenylyl-sulfate reductase gene (glutathione), denoted as *APR*, for the same function. Finally, APS was oxidized to sulfate by the *sat*-coding enzymes, which was found in Ascomycota, Basidiomycota, Chytridiomycota, and Mucoromycota. The genes encoding taurine dioxygenase (*tauD*) for converting taurine to sulfite, were found in Basidiomycota and Ascomycota, like Agaricomycetes, Dothideomycetes and Eurotiomycetes. The genes encoding alkanesulfonate monooxygenase (*ssuD*) for converting alkanesulfonate to sulfite were found in Leotiomycetes (belonging to Ascomycota). These enzymes would produce sulfite from an organic sulfur substrate, which in turn would be fed into the sulfur metabolic pathways.

The hadal sediment fungi were also found to possess sox genes, indicating their potential for thiosulfate/sulfide oxidization. For example, Endogonomycetes of Mucoromycota contained soxC encoding sulfane dehydrogenase subunits for oxidizing thiosulfate to sulfate. Malasseziomycetes of Basidiomycota and Sordariomycetes of Ascomycota were found to have SQOR (eukaryotic sulfide quinone oxidoreductase) genes that are involved in the first step of hydrogen sulfide metabolism to produce sulfane sulfur metabolites. A previous study reported that fungi could feed sulfate to sulfate-reducing bacteria (SRB)[51]. Interestingly, we also found SRB, e.g. Desulfobacterales and Desulfuromonadales (within the class Deltaproteobacteria), and Nitrospirae (class) in the Challenger Deep sediment, which corroborates the evidence for the existence of sulfide oxidation reactions and the roles that fungi that play in sulfur cycling in the hadal sediment habitat. In addition, the hydrogen sulfide metabolisms possessed by the hadal sediment fungi would counter the accumulation of sulfide (H_2_S, HS^-^ and S^2-^) that was generated by SRB in the anaerobic environment. Thus, the discovery of the critical metabolic genes in sulfur metabolism implicated the important role of hadal fungal community in sulfur cycling and energy transformation in the trench environments.

### Metavirome in the Challenger Deep sediment habitat

The existence and composition of viral community in the hadal sediment habitat were an open question remaining to be answered. Our deep metagenomics approach offered a new opportunity to look into the viral components for the deepest biosphere. Viral sequence reads and their taxonomic affiliations were determined using Kaiju by mapping to the viral references from NCBI-nr database. A total of fifteen major viral families were identified in the Challenger Deep sediment (Fig. 4A). dsDNA viruses were found most frequently, which were mainly affiliated with the order Caudovirales, also known as the tailed bacteriophages [52]. The predominant family was Myoviridae, accounting for 24.62% of the total viral sequences, followed by Siphoviridae (19.70%) and Podoviridae (9.40%). Notably, we detected a relatively high abundance of giant viruses in the Challenger Deep sediment, such as Mimiviridae (4.16%) and Phycodnaviridae (1.06%) (Fig. 4A, red star). Giant viruses were often missed or underestimated in early studies due to the technique used to capture viral particles via filtering [53]. Our approach is indifferent to the viral components of different particle sizes.

**Fig. 4.**
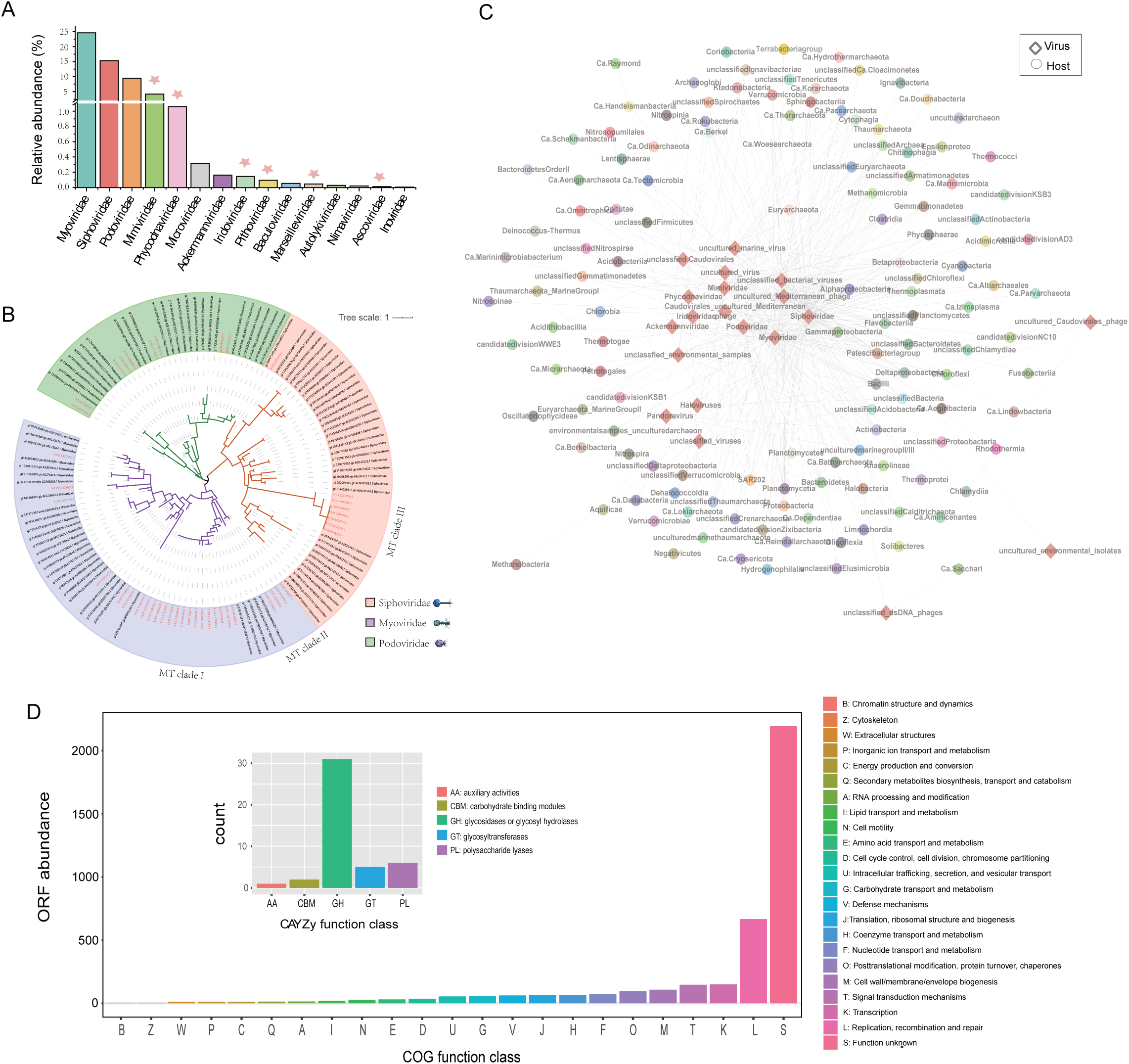
Diversity of metavirome, virus-host association, and viral genome annotation. (A) The relative abundance of dominant virus groups within metavirome. (B) Phylogenetic analysis of Caudovirales based on TerL using the maximum likelihood algorithm. Reference viral sequences from NCBI are colored in black. Scale bar, one amino acid substitution per site. The same tree with detailed labeling is provided in Fig. S3 (Additional file 2: Fig. S3). (C) Visualization of the virus-host association network. Diamonds and circles denote marine viruses and microbial hosts, respectively. Association is summarized between a virus family and a microbe class, represented by a linked grey line. (D) Annotation of viral genomes. COG annotation was performed using eggNOG-mapper [90]. AMGs related to carbohydrate metabolism are illustrated in the inner panel.

To investigate the diversity and evolution among the members of Caudovirales found in the Challenger Deep sediment, phylogenetic analysis was performed on the viral contigs, using a marker gene coding the terminase large subunit (TerL) [54]. Interestingly, for the two families, Myoviridae and Siphoviridae, there were separate branche(s) forming within each of them (Fig. 4B). They were tentatively named MT clade I and MT clade II for Myoviridae, and MT clade III for Siphoviridae. The new diversity of Caudovirales in the Challenger Deep sediment may reflect the differentiation of viral niche to microbial hosts living in the hadal habitat of high-hydrostatic pressure and oligotrophy.

#### Virus-host association network

To infer virus and microbial host interactions, we took the use of a combination of different methods as described [55]. Possible virus-host interactions were summarized for each viral family. As a result, members from 131 bacteria and archaea classes were associated with 20 viral families (Fig. 4C and Additional file 1: Table S9). The association network revealed many one-to-many relationships between viruses and hosts, and vice versa. Siphoviridae was connected with the largest number of hosts that belong to 95 classes. Notably, many of the hosts were uncultured microbes that were only inferred from their presence in metagenome sequences.

Unlike in epipelagic and mesopelagic ocean waters, where the most frequent hosts were Cyanobacteria and Alphaproteobacteria (mainly SAR11 [52]), the most frequent hosts in the Challenger Deep sediment were Firmicutes (mainly Bacilli) and Bacteroidetes, followed by Euryarchaeota (including uncultured marine group II/III and Diaforarchaea), unclassified Chloroflexi, and Alphaproteobacteria. A previous study reported that the dsDNA virus T7virus (belonging to Podoviridae) could infect Deltaproteobacteria, Gammaproteobacteria, Alphaproteobacteria, Firmicutes, and Cyanobacteria [55]. Our results indicated that the T7virus from the Challenger Deep sediment was also associated with an archaeal host, e.g. the Diaforarchaea group of Euryarchaeota.

#### Functional viromics

To investigate the gene contents and functions of the hadal sediment viruses, we annotated their contigs using the Clusters of Orthologous Groups (COGs) database [56]. While a large proportion of their ORFs had unknown functions, high-abundance genes were found in the COG categories of ‘replication, recombination and repair’, ‘transcription’, ‘signal transduction mechanism’, ‘cell wall/membrane/envelope biogenesis’, etc (Fig. 4D). Besides the genes for basic viral functions, a special group of virus-encoded genes can modulate the activities of the hosts upon infection. These so-called auxiliary metabolic genes (AMGs) were a means for viruses to manipulate host metabolism, like sulfur and nitrogen cycling.

We identified 40 putative AMGs from the Challenger Deep sediment viruses, having roles in carbon, sulfur or nitrogen metabolism (Additional file 1: Table S10). Among them, auxiliary carbohydrate metabolic genes were the most frequent. By searching against the CAZymes database, 45 carbohydrate metabolism-related genes were identified, which included 31 GHs (glycosidases or glycosyl hydrolases), six PLs (polysaccharide lyases), five GTs (glycosyltransferases), two CBM (carbohydrate binding modules), and one AA (auxiliary activities). The presence of frequent auxiliary carbohydrate metabolic genes provided evidence that the hadal sediment viruses may heavily manipulate carbohydrate metabolism of hosts, especially promoting carbohydrate degradation on hosts in an oligotrophic habitat. AMGs for carbohydrate metabolism were also the largest group in other marine virome dataset, like the Tara Oceans Viromes [57], which concentrated, however, on different pathways, like galactose metabolism, Glycosyltransferases, etc., reflecting the differentiated viral niche to hosts in dramatically different habitats.

For nitrogen metabolism, *NirK* that encodes nitrite reductase [58] was found in the hadal viruses, suggesting possible roles to enhance host nitrite reduction pathway and release of nitric oxide into hadal sediment. For sulfur cycling, the *CysN/NodQ* gene encoding ATP sulphurylase [59] that reduces sulfate to produce 3’-phosphoadenosine-5’-phosphosulfate (PAPS), the first step of assimilatory sulfate reduction, was found in the hadal viruses. Notably, both *NirK* and *CysN/NodQ* are found for the first time among AMGs for ocean viruses.

### Large-scale isolation of microbes from the Challenger Deep sediment habitat

To catalog and characterize microbes in the hadal sediment habitat, an intensive effort was made for the isolation of microbes from the Challenger Deep sediment. We adopted a protocol previously developed for the isolation of environmental microorganisms, which used serial dilutions of samples to allow the growth of ‘disadvantaged’ microbes that otherwise would not be isolated [60]. To broaden the diversity of microbial isolates, we employed 24 different media and combined them with various culture conditions (Additional file 1: Table S11). Notably the experimental conditions would enable facultative anaerobes but not strict anaerobes. As a result, more than two thousand individual isolates were obtained, for which 1089 were completed for 16S rRNA gene (for prokaryotes) or ITS amplicon (for eukaryotes) sequencing (Additional file 1: Table S12 and S13).

#### Bacterial isolates

The taxonomy of 1070 bacterial isolates was assigned based on 16S rRNA gene sequences using SILVA database (release 123) [61]. The bacterial isolates belonged to four phyla, i.e. Proteobacteria, Bacteroidetes, Actinobacteria, and Firmicutes, which were further categorized into 7 classes, 18 orders, 25 families, and 40 genera (Additional file 2: Fig. S2 and Additional file 1: Table S12). They matched the microbes represented by sequences in the metagenome. Halomonas, Pseudoalteromonas, and Idiomarina were the top genus, accounting for 32.77%, 22.23%, and 9.92% of the isolates, respectively (Fig. 5A). Nineteen bacterial isolates from five classes were candidates for new species, which had a 16S rRNA gene sequence with <=97% identity to their closest references (Additional file 1: Table S14)[62].

**Fig. 5.**
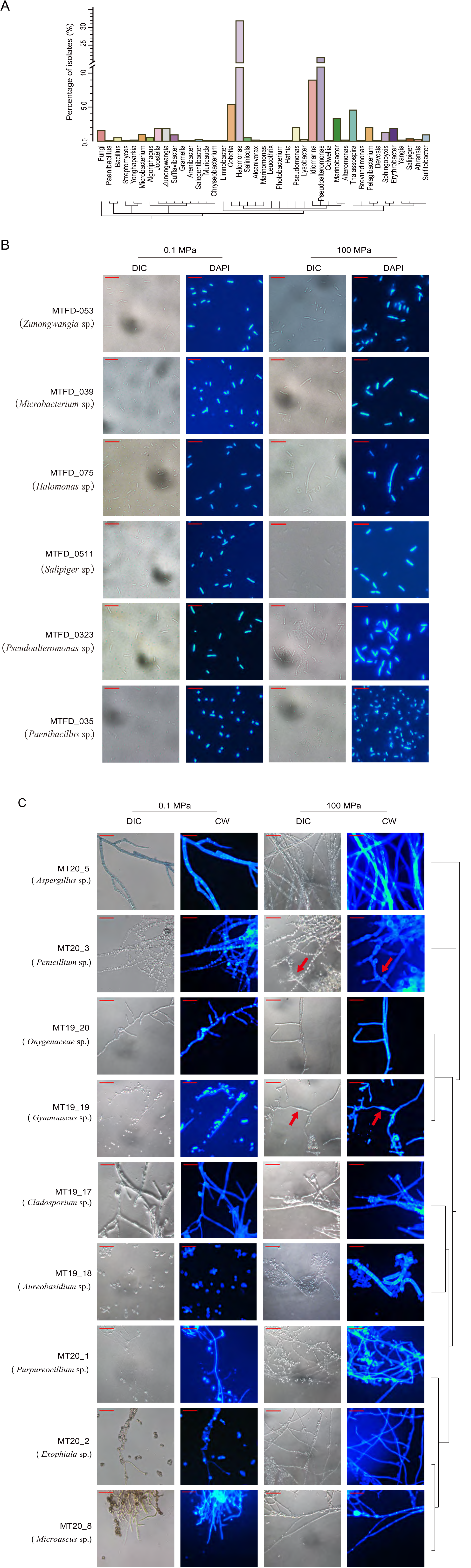
Bacterial and fungal isolates from the Challenger Deep sediment habitat. (A) Phylogeny and percentage of bacterial and fungal isolates (Additional file 1: Table S12 and S13). (B) Visualization of representative bacterial isolates from four different phyla. Cells were cultured under 0.1 and 100 MPa, respectively. Scale bars, 10 μM; DIC, images taken with a DIC microscope; DAPI, cells stained with DAPI and observed with an epifluorescence microscope. (C) Visualization of representative fungal isolates from nine genera. Scale bars, 30 μM; CW, cells stained with Calcofluor White and observed with an epifluorescence microscope. The phylogeny of fungal isolates is shown to the right. Red arrows indicate swollen hypha (MT19_19) or swollen conidia (MT20_3).

We characterized representative isolates from each of the four phyla, i.e. MTFD_053 (*Zunongwangia* sp.), MTFD_039 (*Microbacterium* sp.), MTFD_075 (*Halomonas* sp.), MTFD_0511 (*Salipiger* sp.), MTFD_0323 (*Pseudoalteromonas* sp), and MTFD_035 (*Paenibacillus* sp.), which were visualized after incubation at elevated pressure (100 MPa) for seven days. Some morphologic changes were observed with MTFD_039, MTFD_075, MTFD_0323 and MTFD_0511, when compared between their cells cultured at 0.1 and 100 MPa (Fig. 5B). Cases were found that sister cells were connected after cell divisions under culture at elevated pressure. These isolates exemplified the small fraction of culturable microbes in contrast to the community background in the Challenger Deep sediment illustrated by the MAG data, which complemented the lack of taxonomic details from the metagenomic analysis. The results suggested that they were piezotolerant, which are likely derived from the microbes that descended from the water column, contributing to the diversity and metabolic functions of the hadal sediment microbiome.

#### Microeukaryotic isolates

Nineteen microbial eukaryotes (fungi) were isolated from the Challenger Deep sediment (Additional file 1: Table S13 and Additional file 2: Fig. S4). Their taxonomy was assigned based on their ITS sequences using the NCBI ITS database. They belonged to nine fungal genera, namely Aspergillus, Penicillium, Acremonium, Microascus, Exophiala, Cladosporium, Gymnoascus, Purpureocillium, and Aureobasidium. They matched the eukaryotic microbes represented by sequences in the metagenome.

Representative isolates from each genus were further characterized after incubation at 0.1 and 100 MPa for fourteen days. MT19_18 (*Aureobasidium* sp.) tended to form long pseudomycelia under 100 MPa (Fig. 5C). Some elongated cells contained multiple nuclei but did not have a diaphragm separating them. The filamentous fungi, i.e. MT20_2 (*Exophiala* sp.), MT20_5 (*Aspergillus* sp.), MT20_3 (*Penicillium* sp.), MT19-20 (*Onygenaceae* sp.), MT19_19 (*Gymnoascus* sp.), MT19_17 (*Cladosporium* sp.), MT20_1 (*Purpureocillium* sp.), and MT20_8 (*Microascus* sp.), have some morphologic variations induced by elevated pressure, such as hyphal swelling (MT19_19) and swollen conidia (MT20_3) (Fig. 5C). Conidiophore appeared normal for MT19_17 under 100 MPa compared to that under 0.1 MPa. MT20_1 (*Purpureocillium* sp.), MT19_19 (*Gymnoascus* sp.), and MT20_8 (*Microascus* sp.) may represent new species specialized in the Challenger Deep habitat, whose ITS sequences diverged from their closest known species (Additional file 1: Table S13).

## Conclusions

We adopted a two-fold strategy to investigate the microbial community structure and functions in the Challenger Deep sediment biosphere. The deep metagenomics approach reconstructed 178 MAGs from the deepest habitat, and revealed the full biosphere structure and novel biodiversity, particularly enabling and extending our study to include microeukaryotic and viral components. The largest MAG set reconstructed for the hadal microbiome, established versatile community functions marked by unexpected enrichment and prevalence of mixotrophy and facultative anaerobic metabolism. On the other hand, fungi as heterotrophs had broad metabolic potentials in carbon, nitrogen, and sulfur metabolism, particularly the capability of dissimilatory nitrate reduction and sulfate reduction that had not been reported for deep ocean microeukaryotes. These findings implicated the possible roles of hadal fungi in the biogeochemical processes of the hadal trench environment. The viral components evolved with new diversities and carried AMGs important for modulating hosts’ metabolic functions, which illustrated the differentiation of viral niche to microbial hosts in the hadal habitat.

The large-scale cultivation approach obtained more than 2000 isolates from the Challenger Deep sediment, and cataloged 1070 bacteria and 19 fungi by 16S rRNA gene or ITS amplicon sequencing. Many of them were likely new species specialized in the Challenger Deep habitat, and showed morphological variations under elevated pressure (100 MPa). These isolates represent a small fraction of the diversity in the Challenger Deep sediment habitat based on the deep metagenomic data. They would serve as model organisms and provide new opportunities to study and understand the physiology of piezotolerant microbes.

## Materials and Methods

### Sample collection and geochemical measurements

The one intact sediment core was obtained from the seafloor in the Challenger Deep, the Mariana Trench (142°21.7806’E, 11°25.8493’N, 10840 meter) during the cruise from December 2016 to January 2017 onboard the ship ZhangJian. The sediment core was collected using a box corer (with a base area of 400 cm^2^ and a height of 25 cm) attached to the hadal lander-II, developed by Shanghai Engineering Research Center of Hadal Science and Technology, Shanghai Ocean University. After the lander reached the seafloor, the sampling chamber was slowly driven into the subseafloor until it reached around 21 cm below the sediment surface. A lid was then released to seal the box corer, before the lander was recovered. Once the lander surfaced and box corer was recovered and examined for integrity. Only one intact sediment core with well-preserved sediment stratification was used in this study. The sediment core was immediately subsampled by inserting polycarbonate tubes to preserve the segments. The samples were either frozen at -80 °C before stored at -20 °C or stored at 4 °C for culturing and isolation of microbes. Note, the temperature of the recovered core was not obtained. For reference, the recorded environment temperature at the time was 2-4 °C at the ocean floor, and 27-28 °C at the ocean surface.

Total organic carbon (TOC), total nitrogen (TN), and carbon and nitrogen isotopic compositions of particulate organic matter were measured by high-temperature combustion on a Vario Pyro Cube elemental analyzer connected to an Isoprime 100 continuous flow isotope ratio mass spectrometer. All samples were pre-treated with 10% HCl to remove inorganic carbon. Carbon and nitrogen isotope ratios are expressed in the delta notation (δ^13^C and δ^15^N) relative to V-PDB and atmospheric nitrogen. The average standard deviation of each measurement, determined by replicate analyses of the same sample, was ±0.02% for TOC, ±0.006% for TN, ±0.2‰ for δ^13^C, and ±0.3‰ for δ^15^N.

For the measurement of major and minor elements, approximately 40 mg of dry sediment powder was weighed into a Teflon beaker and dissolved in super-pure HF (0.2 ml), HNO_3_ (0.8 ml), and HCl (0.1 mL). The beaker was then sealed and heated on a hot plate at 185 °C for 36 h. After cooling, the solution was evaporated at 120 °C to dryness. The residue was re-dissolved by adding 2 ml HNO_3_ (super-pure) and 3 ml of deionized water at 135 °C in an airtight beaker for 8 h. The final solution was diluted to 50 ml with 3% HNO_3_. Blanks, duplicate samples, and several certified reference materials (GSR-1, OU-6, 1633-a, GXR-2, GXR-5) were also prepared using the same procedure. Major and minor elements were determined using a Thermo-Fisher iCAP6300 ICP-OES and a Perkin-Elmer ELAN 6000 ICP-MS, respectively. The analytical precision was 5% for major and 10% for minor elements.

Measurements of NO_2_^-^, NO_3_^-^, NH_4_^+^, and PO_4_ were performed using a QuAAtro autoanalyzer (Seal Analytical) with a detection limit of 1 μM and a precision of 2%. Sulfate (SO_4_^2-^) was measured by a Dionex ICS-5000^+^ ion chromatograph with a detection limit of 10 μM and a precision of 2%.

### Nucleic acid extraction and metagenomic sequencing

DNA was extracted from 10 g of sediment sample for each experiment replicate, using the MoBio PowerSoil DNA Isolation kit (MO BIO Laboratories, USA) according to the manufacturer’s protocol. DNA concentrations of the extract were measured with a Qubit fluorometer. For each library construction, about 100 ng DNA was fragmented with Covaris S2 (Covaris, USA) and was used to construct metagenomic DNA library with NEXTflex™ DNA Sequencing Kit compatible with the Biomek® FXp (Bio Scientific, USA). Notably PCR-amplification was limited to 12 cycles for each Illumina library. The quality of DNA library was examined by Agilent Bioanalyzer 2100 (Agilent, USA) with DNA 12000 Kit. Paired-end Illumina sequencing (2 x 150 bp) was performed for each metagenomic library on Hiseq Xten instruments (Illumina).

### Metagenomic assembly, mapping, and binning

Raw sequence data were cleaned by removing low-quality sequence reads, artificial sequence reads and contaminated sequences with the following steps: 1) Reads with average quality score less than Q20 and a length < 30 bp, or with adapter sequences, were removed with trimmomatic (version 0.38) [63]; 2) Reads with contaminated sequences, like plasmid sequences, human sequences, etc, were removed with bowtie2 (v2.3.4.1) [64]; 3) Duplicated reads generated by PCR amplification were removed for metagenomic assembly using fastuniq [65]. The metagenome was assembled with clean reads using MEGAHIT (v1.1.3) [66] with the following parameters: --k-list 21, 29, 39, 59, 79, 99, 119, 127, 139. Read coverage for contigs was determined by mapping sequencing reads to contigs using bowtie2 with default parameters.

Metagenomic binning and refinement were performed on contigs using a combination of different tools, including CONCOCT [67], Metabat [68], DAS Tool (v1.0) [69], and metaWRAP [70]. Briefly, initial binning was conducted with CONCOCT and Metabat based on tetranucleotide frequencies of contigs and coverage depth as covariance, using a contig length cutoff of 1 or 1.5 kb, respectively, for which 310 and 400 bins were recovered. Results from the two different binning tools were then combined and merged using the DAS Tool. The resulting bins were refined and consolidated into the final bin sets using metaWRAP’s Bin_refinement module with options: -c 50 -x 10. As a result, 418 bins were recovered across the entire quality spectrum. The quality of the binning results was evaluated by estimating the completeness and contamination scores using CheckM (v1.0.5) lineage_wf tool [32]. A cutoff of 50% completeness and 10% contamination was used to filter and obtain quality genomes (MAGs).

### Taxonomic classification of sequences and determination of relative sequence abundance

The method for taxonomic assignment of sequence reads and contigs was adapted from the previous work using Kaiju [20]. Briefly, the NCBI-nr database compiled in a GORG-Tropics format was downloaded, which included reference sequences from archaea, bacteria, viruses, and microbial eukaryotes. The assignment of the sequence reads or contigs was conducted using the NCBI-nr database and Kaiju (v1.7.0) [71] in Greedy-5 mode with the default options. Sequences were assigned with the taxonomic id and functional annotations based on mapped references. Their taxonomic lineage information was then obtained using Kaiju’s tool ‘addTaxonNames’. For sequence reads assigned with taxonomic id, the relative sequence abundance of a phylum, class, order, etc. was estimated by summarizing the total number of assigned sequence reads for a given category and dividing it with the total number of assigned reads.

### Phylogenetic analysis and taxonomy assignments of the metagenome-assembled genomes (MAGs)

The sequences of 43 conserved proteins from previous work [32] were retrieved from the metagenome-assembled genomes (MAGs), and multi-aligned using the MUSCLE program (v3.8.31) [72]. The alignments were trimmed to remove gaps and poorly aligned regions using TrimAL with the options -gt 0.95 -cons 50 [73]. The ‘cleaned’ alignment datasets for the 43 conserved proteins were then concatenated, and were subsequently used for constructing a phylogenetic tree using the maximum likelihood algorithm. The RAxML program (v8.1.24) [74] was employed with the options “-f a -n boot -m PROTGAMMALG -c 4 -e 0.001 -# 1000”. To visualize the phylogenetic tree topology, the Newick files were processed using iTOL online tool (v4) [75].

Initial taxonomy assignments of MAGs were provided from the evaluation of MAGs with CheckM, which were often short with taxonomic information for the order, family, or genus. The MAGs were further classified using CAT and GTDB-Tk programs [33, 76]. CAT assigns the taxonomy of MAGs based on the BLAST results of the nr database using the lowest common ancestor (LCA) algorithm [76]. On the other hand, GTDB-Tk provides taxonomic classifications with the new rank-normalized GTDB taxonomy, by using two criteria, relative evolutionary divergence (RED) and average nucleotide identity (ANI) for establishing taxonomic ranks [27, 33]. The taxonomy assignment for MAGs were determined to the deepest taxonomic levels by combining the results from both CAT and GTDB-Tk analyses. A few inconsistent assignments between the two programs were manually resolved based on information of their nearest reference neighbors.

### Annotation of contigs and metabolic pathway analysis

Genes were predicted for contigs / metagenome-assembled genomes (MAGs) using prodigal (v2.6.2) with -p meta parameters [77]. Putative genes were then annotated with KEGG Automatic Annotation Server (KAAS) using the GHOSTX tool with the ‘custom genome dataset’ and ‘BBH’ options, by uploading their predicted amino acid sequences to KAAS [78]. Furthermore, putative genes were annotated using blastp program with default options against several different databases, like hmmer, the curated database of Anaerobic Hydrocarbon Degradation Genes (‘AnHyDeg’), and MEROPS database [42]. To identify genes encoding carbohydrate degradation enzymes, the dbcan tool was used to search the Carbohydrate-Active enzymes (CAZYmes) database with an e-value cutoff of 1e-5 [79]. To reconstruct metabolic pathways for a draft genome (MAG), its predicted genes were annotated and pathway results were summarized using the KEGG server. To illustrate metabolic activity for certain pathway(s) within a taxonomic unit (e.g. phylum, class, order, etc) or a phylogenetic cluster, all genomes within the specific taxonomic unit/cluster were summarized, based on their taxonomic or phylogenetic assignments.

### Identification of viral contigs and phylogenic analysis

Viral contigs in the metagenome assembly were identified using Kaiju (v1.7.0). For the phylogenic analysis of Caudovirales, the TerL gene [80] was identified for each contig using blastp with a threshold e-value of 10^-5^, minimum identity of 50%, and minimum coverage of 30%. Identified TerL gene sequences were retrieved from contigs and aligned using the Mafft program (version 7.407) [81]. The alignments were trimmed to remove gaps and ambiguously aligned regions before the phylogenetic tree was constructed using the Fasttree program [82], and manually formatted and visualized using iTOL.

### Virus - host association and viral function analysis

Viruses and their possible hosts from the hadal sediment environment were inferred using the previously developed methods [55]. Briefly, virus-host association was predicted using three different means: 1) Virus contig - host genome similarity analysis: All identified viral contigs from the sediment of the Challenger Deep were compared to the archaeal and bacterial contigs in the same metagenome, using blastn with a threshold of 50 for bit score and 0.001 for E-value. 2) Virus contig - host CRISPR spacer match analysis: CRISPR spacers were predicted for all microbial contigs using MetaCRT [83]. Association between a virus and its hosts was predicted when a host CRISPR spacer was found to match a virus contig. 3) Nucleotide composition comparison between virus contigs and host genomes. The tetranucleotide-frequency vectors and mean absolute error between the vectors for each virus-host pair were computed. A viral contig was assigned to the closest microbial host whose genome having the lowest mean absolute error (d) to the viral contig if d < 0.0015.

For functional viromics analysis, COG annotation of viral contigs was performed using eggNOG-mapper (http://eggnog-mapper.embl.de/) with a threshold of 10^−5^ Viral ORFs annotated as the CAZymes family genes in eggNOG-mapper annotation results were considered as auxiliary metabolic genes (AMG) related to carbohydrate metabolism [84]. Further, additional viral ORFs related to sulfur and nitrogen metabolism (determined by KEGG kos) in the eggNOG-mapper annotation results were also considered as AMGs.

### Large-scale cultivation and identification of the hadal sediment microbes

The cultivation and isolation of the microbes from the Challenger Deep sediment were adopted from previous studies as described [60, 85]. To reduce their exposure to oxygen, the sediment sub-samples were sealed in air-tight bags with residue air removed before they were stored at 4 °C. A series of dilutions, e.g. 10^-1^, 10^-2^, 10^-3^, 10^-4^, and 10^-5^, were made to the sediment samples in 96-well plates using various culture media (Additional file 1: Table S11). The high ratio dilutions, e.g. 10^-3^, 10^-4^, 10^-5^, were then plated out in 90-mm plates with solid culture media. The dilution and inoculation were performed under aerobic conditions in the laboratory. The plates were incubated under various conditions, i.e. 4°C or 28°C, aerobic or anaerobic, with or without antibiotic, etc, for up to one month. For anaerobic culture, the agar media plates were first treated with AnaeroPack (Mitsubishi Gas Chemical Company, Inc., Tokyo) to remove oxygen. For large-scale isolation, a combination of 24 different media (modified from nine base-types) with various culture conditions were used for cultivation (Additional file 1: Table S11). Single colonies emerging at different time points were picked to grow in liquid media for cryopreservation and further analysis. Notably some single colonies emerging at late stages were collected from the highest dilution plates, which would have been overlaid by faster-growing colonies in the low-dilution plates.

The identity of the single colonies was analyzed by 16S rRNA gene (for prokaryotes) or ITS gene (for eukaryotes) sequencing. The primer sets, 8F/805R (8F-5’AGAGTTTGATCCTGGCTCAG; 805R-5’GACTACCAGGGTATCTAATC) (targeting V1-V4 regions) and 8F/1492R (8F-5’AGAGTTTGATCMTGGCTCAG; 1492R-5’GGTTACCTTGTTACGACTT) (targeting V1-V9 regions) were used for 16S rRNA gene tag PCR-amplification and sequencing[86, 87]. They generated the PCR-amplified tag with length of 805 and 1492 bp, respectively. The primer set, ITS-F/ITS-R (ITS-F-5’GGAAGTAAAAGTCGTAACAAGG; ITS-R-5’TCCTCCGCTTATTGATATGC) were used for ITS gene tag PCR-amplification and sequencing. It generated the PCR-amplified tag with length of of 578 bp. Identification of the sequenced single colony was conducted by searching 16S ribosomal RNA sequence database, and Internal transcribed spacer region (ITS) database at NCBI (https://www.ncbi.nlm.nih.gov/) using blastn.

### Culture of microbes under elevated hydrostatic pressure and microscopic imaging

Single colony was picked from solid media to inoculate 2-ml liquid culture media in a sterile 15-ml glass tube, and was incubated on a rotary shaker (180 rpm) at 28 °C overnight for bacteria or for three days for fungi to produce seed broth. For single-cell microbes, cultures were diluted to O.D.600 = ∼0.01 in appropriate media. For filamentous fungi, mycelia in the seed broth were fragmented with sterilized scissors (if necessary) before the cultures were diluted (1/15 ∼ 1/60 depending on fungal biomass) in the same media. The seeded cultures were transferred to 2 ml sterile plastic syringes before they were placed inside pressure vessels (Model: FY2016108; Feiyu Petroleum Technology Development Co., Ltd., Nantong, China). Hydrostatic pressure of 100 MPa was applied at room temperature to bacterial cultures for seven days, or to fungal cultures for fourteen days, respectively. After high-pressure incubation, the syringes were taken out of pressure vessels and visually examined for physical integrity. The cultures were re-plated on solid media to test viability for growth or were examined by microscopy.

For microscopic imaging of bacteria and fungi, an epifluorescence microscope (Nikon Ecllipse 80i) and a digital camera (Nikon DS-Ri1; software: NIS-Elements F Ver4.30.01) attached to the microscope were used. For bacterial culture, cells were harvested by centrifugation. Pellets were washed twice with PBS buffer, and stained with DAPI (4’,6-diamidino-2-phenylindole) for 10 minutes in the darkness [88]. Stained cellular suspension (2.5 μL) was spread onto a class microscope slide, and covered with a cover slide with excess liquid removed. Slides were imaged at x100 magnification (oil immersion len) for Differential Interference Contrast (DIC) micrograph and epifluorescent micrograph (excitation wavelength 300-380 nm), respectively. For fungal culture, 10 μL fungal culture was mixed with 10 μL Calcofluor White solution (1 mg/mL) on a glass slide for 3 min at room temperature [89]. Fungal hyphae with or without Calcofluor staining were imaged at x40 magnification for DIC micrograph and epifluorescent micrograph, respectively.

## Availability of data and materials

The metagenomic sequence data have been deposited in the NCBI Sequence Read Archive (SRA) under accession PRJNA723166 [91]. The 16s rRNA/ITS tag sequences have been deposited to NCBI GenBank (Accession numbers listed in Tables S12 and S13).

## Acknowledgments

We thank the reviewers for their constructive comments which have greatly assisted us in improving this manuscript.

## Ethics declarations

### Ethics approval and consent to participate

Not applicable

### Consent for publication

Not applicable.

### Author contributions

X.L. conceived the project and wrote the manuscript. J.F. and P.N. advised on the study, and directed experiments for bacterial isolation and high-hydrostatic cultivation. P.C., H.Z., and J.L. conducted metagenomics experiments, analyzed data, and prepared the manuscript. Y.H., M.Z., Z.X., T.J., and W.D. performed bacterial cultivation and isolation. Y.W., J.C., and M.L. conducted geochemistry experiments and data collection.

## Supplementary information

### Additional file 1: Supplementary Tables S1-S16

**Table S1.** Geochemical characterization of the sediment samples from Challenger Deep.

**Table S2.** Concentrations of nutrient ions NO_3_^-^, NO_2_^-^, PO_4_^-^, NH_4_^+^, and SO_4_^2-^ in porewater of the samples.

**Table S3.** Dissolved major elements in the sediment samples.

**Table S4.** Dissolved trace elements in the sediment samples.

**Table S5.** Deep metagenomic sequencing on the Challenger Deep sediment samples and assembly.

**Table S6.** Taxonomic assignment and characteristics of the 178 MAGs.

**Table S7.** Functional analysis of archaeal and bacterial MAGs. The key genes of each metabolic pathways for MAG annotation are listed on the top. Presence/absence of genes are listed as: Presence: 1 (green); Absence: a (no color). The module completeness of each metabolic pathway is the percentage of the encoded key genes in the corresponding pathway (e.g. module completeness value of 25 means the MAG contained one of the four key genes in the pathway).

**Table S8.** The metabolic characteristics of the six major the Challenger Deep Fungi groups, and detailed information of the carbohydrate-active enzyme family (CAZymes) and peptidase family genes detected in major fungal groups.

**Table S9.** List of host prediction for the Challenger Deep virome. For each prediction, the type of signal (blastn, CRISPR, tetranucleotide composition), the host sequence used for the prediction alongside its affiliation, and the strength of the prediction (length of the blastn match, number of mismatches in the CRISPR spacer, and distance between viral and host tetranucleotide frequencies vectors) are indicated.

**Table S10.** List of eggNOG-mapper annotations for the Challenger Deep virome.

**Table S11.** Twenty-four different media modified from several base-type that favored the growth of different bacteria or fungi.

**Table S12.** The taxonomic assignment of the 1070 bacterial isolates. Their taxonomy was assigned based on 16S rRNA gene sequences using NCBI rRNA/ITS databases.

**Table S13.** Characterization of nineteen sequenced Fungi.

**Table S14.** Nineteen bacterial isolates are considered as new species, which have 16S rRNA gene sequences with <=97% identity to closest reference.

**Table S15.** Composition of microbial communities with other Mariana trench sites.

**Table S16.** The relative abundance comparison of dominant microbial groups in Mariana Trench sediment with other hadal sediment samples.

### Additional file 2: Supplementary Figs. S1-S4

**Fig. S1.** Phylogenetic analyses of bin.150 and some other Candidatus Diapherotrites archaeon reference genomes downloaded from the NCBI database. A FastTree approximate maximum-likelihood phylogenetic tree was built using 15 conserved ribosomal protein.

**Fig. S2.** Phylogenetic analyses of 1070 bacterial isolates from the hadal sediment based on the 16S rRNA sequence with the FastTree. The color bar represents the order level.

**Fig. S3.** Phylogenetic analysis of Caudovirales based on TerL using the maximum likelihood algorithm. Reference viral sequences from NCBI are colored in black. Scale bar, one amino acid substitution per site.

**Fig. S4.** Culture of fungi under elevated hydrostatic pressure. A) High-pressure cultivation system. B) Nineteen fungi isolated from the Challenger Deep sediment, which was tested to be viable to grow on plates after subjecting to high hydrostatic pressure (100 MPa) for fourteen days. Isolate names are listed at the bottom.

**Additional file 3:** Analysis of MAGs on potential for different types of fermentation.

