## Additional file 1: Table S1 for "Revealing the full biosphere structure and versatile metabolic functions in the deepest ocean sediment of the Challenger Deep"

**Additional file 1: Table S1.** Geochemical characterization of the sediment samples from the Challenger Deep.

| **Sample ID** | **MT-1** | **MT-2** | **MT-3** |
| --- | --- | --- | --- |
| Water Depth (m) | 10840 | 10840 | 10840 |
| Sediment core depth (cm) | 0-5 | 5-10 | 10-14 |
| Temperature (℃) | 2.5 | 2.5 | 2.5 |
| Salinity (PSU) | 34.6810 | 34.6810 | 34.6810 |
| TOC (wt%) | 0.55 | 0.49 | 0.52 |
| TN (wt%) | 0.06 | 0.05 | 0.06 |
| TOC/TN (mol/mol) | 10.07 | 10.86 | 9.91 |
| δ^13^C (‰) | -21.41 | -21.64 | -21.53 |
| δ^15^N (‰) | 6.42 | 5.42 | 6.69 |

**Note:**

Total organic carbon (TOC), total nitrogen (TN), and carbon and nitrogen isotopic compositions of particulate organic matter were measured by high-temperature combustion on a Vario Pyro Cube elemental analyzer connected to an Isoprime 100 continuous flow isotope ratio mass spectrometer. All samples were pre-treated with 10% HCl to remove inorganic carbon. Carbon and nitrogen isotope ratios are expressed in the delta notation (δ^13^C and δ^15^N) relative to V-PDB and atmospheric nitrogen. The average standard deviation of each measurement, determined by replicate analyses of the same sample, was ±0.02% for TOC, ±0.006% for TN, ±0.2‰ for δ^13^C, and ±0.3‰ for δ^15^N.
