## Additional file 1: Table S2 for "Revealing the full biosphere structure and versatile metabolic functions in the deepest ocean sediment of the Challenger Deep"

**Additional file 1: Table S2**. Concentrations of nutrient ions NO_3_^-^, NO_2_^-^, PO_4_^-^, NH_4_^+^, and SO_4_^2-^ in porewater of the samples.

| **Sample ID** | **MT-1** | **MT-2** | **MT-3** |
| --- | --- | --- | --- |
| **Depth (cm)** | **0-5** | **5-10** | **10-14** |
| NO_3_^-^ (μM) | 34.70±0.31 | 32.96±0.46 | 31.49±0.83 |
| NO_2_^-^ (μM) | 0.90±0.58 | 1.84±0.86 | 1.76±1.26 |
| PO_4_^3-^ (μM) | 2.66±0.26 | 3.33±0.58 | 3.87±0.59 |
| NH_4_^+^ (μM) | 0.43±0.36 | 1.23±0.21 | 3.28±0.70 |
| SO_4_^2­-^ (mM) | 28.21±0.20 | 28.18±0.28 | 28.26±0.17 |

**Note:**

The average standard deviation of each measurement, determined by replicate analyses of the same sample. Measurements of NO_2_^-^, NO_3_^-^, NH_4_^+^, and PO_4_^3-^ were performed using a QuAAtro autoanalyzer (Seal Analytical) with a detection limit of 1 μM and a precision of 2%. Sulfate (SO_4_^2-^) was measured by a Dionex ICS-5000^+^ ion chromatograph with a detection limit of 10 μM and a precision of 2%.
