## Additional file 1: Table S3 for "Revealing the full biosphere structure and versatile metabolic functions in the deepest ocean sediment of the Challenger Deep"

**Additional file 1: Table S3.** Dissolved major elements in the sediment samples.

| **Sample ID** | **MT-1** | **MT-2** | **MT-3** |
| --- | --- | --- | --- |
| **Depth (cm)** | **0-5** | **5-10** | **10-15** |
| Na_2_O (wt%) | 1.93 | 1.79 | 1.93 |
| MgO (wt%) | 10.13 | 8.14 | 10.65 |
| Al_2_O_3_ (wt%) | 11.53 | 10.61 | 11.59 |
| P_2_O_5_ (wt%) | 0.20 | 0.19 | 0.19 |
| K_2_O (wt%) | 1.51 | 1.48 | 1.56 |
| CaO (wt%) | 2.29 | 2.10 | 2.42 |
| TiO_2_ (wt%) | 0.57 | 0.54 | 0.60 |
| MnO (wt%) | 0.51 | 0.5 | 0.54 |
| Fe_2_O_3_-T% | 7.75 | 7.06 | 8.31 |

**Note:**

For the measurement of major and minor elements, approximately 40 mg of dry sediment power were weighed into a Teflon beaker and dissolved in super-pure HF (0.2 ml), HNO_3_ (0.8 ml), and HCl (0.1 ml). The beaker was then sealed and heated on a hot plate at 185 °C for 36 h. After cooling, the solution was evaporated at 120 ^o^C to dryness. The residual was re-dissolved by adding 2 ml HNO_3_(super-pure) and 3 ml of deionized water at 135 °C in an airtight beaker for 8 h. The final solution was diluted to 50 ml with 3% HNO_3_. Blanks, duplicate samples, and several certified reference materials (GSR-1, OU-6, 1633-a, GXR-2, GXR-5) were also prepared using the same procedure. Major and minor elements were determined using a Thermo-Fisher iCAP6300 ICP-OES and a Perkin-Elmer ELAN 6000 ICP-MS, respectively. The analytical precision was better than 5% for major and 10% for minor elements.
