## Additional file 1: Table S4 for "Revealing the full biosphere structure and versatile metabolic functions in the deepest ocean sediment of the Challenger Deep"

**Additional file 1: Table S4.** Dissolved trace elements in the sediment samples.

| **Sample** | **MT-1** | **MT-2** | **MT-3** |
| --- | --- | --- | --- |
| **Depth (cm)** | **0-5** | **5-10** | **10-14** |
| Li (μg/g) | 32.03 | 30.65 | 32.67 |
| Be (μg/g) | 0.93 | 0.93 | 0.98 |
| Sc (μg/g) | 17.50 | 16.85 | 17.67 |
| Co (μg/g) | 42.47 | 38.95 | 42.23 |
| Cu (μg/g) | 172.00 | 169.00 | 181.33 |
| Zn (μg/g) | 96.97 | 93.05 | 99.93 |
| Ga (μg/g) | 12.53 | 12.00 | 12.90 |
| Ge (μg/g) | 3.71 | 3.54 | 3.78 |
| As (μg/g) | 8.74 | 8.25 | 9.06 |
| Rb (μg/g) | 39.43 | 39.80 | 41.20 |
| Y (μg/g) | 21.37 | 21.55 | 22.83 |
| Zr (μg/g) | 86.87 | 84.10 | 90.40 |
| Nb (μg/g) | 6.05 | 5.85 | 6.26 |
| Cs (μg/g) | 2.95 | 3.02 | 3.14 |
| La (μg/g) | 17.87 | 18.60 | 18.67 |
| Ce (μg/g) | 35.13 | 35.05 | 37.63 |
| Pr (μg/g) | 4.68 | 4.80 | 5.03 |
| Nd (μg/g) | 19.47 | 20.10 | 20.93 |
| Sm (μg/g) | 4.51 | 4.66 | 4.81 |
| Eu (μg/g) | 1.21 | 1.22 | 1.29 |
| Gd (μg/g) | 4.94 | 5.06 | 5.27 |
| Tb (μg/g) | 0.81 | 0.83 | 0.87 |
| Dy (μg/g) | 4.75 | 4.85 | 5.08 |
| Ho (μg/g) | 1.03 | 1.05 | 1.12 |
| Er (μg/g) | 2.82 | 2.91 | 3.03 |
| Tm (μg/g) | 0.39 | 0.40 | 0.43 |
| Yb (μg/g) | 2.56 | 2.62 | 2.75 |
| Lu (μg/g) | 0.38 | 0.39 | 0.41 |
| Hf (μg/g) | 2.08 | 2.05 | 2.19 |
| Ta (μg/g) | 0.40 | 0.39 | 0.42 |
| Pb (μg/g) | 18.90 | 19.25 | 20.93 |
| Th (μg/g) | 3.44 | 3.60 | 3.73 |
| U (μg/g) | 0.88 | 0.87 | 0.94 |
| Mo (μg/g) | 1.40 | 1.40 | 1.54 |
