## Additional file 1: Table S5 for "Revealing the full biosphere structure and versatile metabolic functions in the deepest ocean sediment of the Challenger Deep"

**Additional file 1:** Table S5. Deep metagenomic sequencing on the Challenger Deep sediment samples and assembly.

|  | **MT-1** | **MT-2** | **MT-3** | | | **Total** | |
| --- | --- | --- | --- | --- | --- | --- | --- |
| Num Illumina Libraries | 3 | 3 | | 5 | | | 11 |
| Raw data (Gb) | 11.7+33.0  +24.11 | 37+26.54  +25.77 | | 26.2+25.6+18.97  +4.73+15.03 | | |  |
| Total Raw data (Gb) | 68.81 | 89.31 | | 90.54 | | | 248.66 |
| Clean data  (Gb) | 6.83+27.68  +21.84 | 32.47+21.61  +23.43 | | 23.11+19.3+17.31  +4.07+13.88 | | |  |
| Total Clean data (Gb) | 56.35 | 77.51 | | 77.67 | | | 211.53 |
|  | Assembly length (bp): | | |  | 6,645,507,297 | | |
|  | Number of contigs: | | |  | 6,208,161 | | |
|  | Average length (bp): | | |  | 1,070 | | |
|  | N50 (bp): | | |  | 1,138 | | |
|  | Max_length (bp): | | |  | 796,916 | | |
|  | Min_length (bp): | | |  | 500 | | |
|  | GC (%): | | |  | 52.28 | | |
