## Additional file 2: Figure S1-S4 for "Revealing the full biosphere structure and versatile metabolic functions in the deepest ocean sediment of the Challenger Deep"

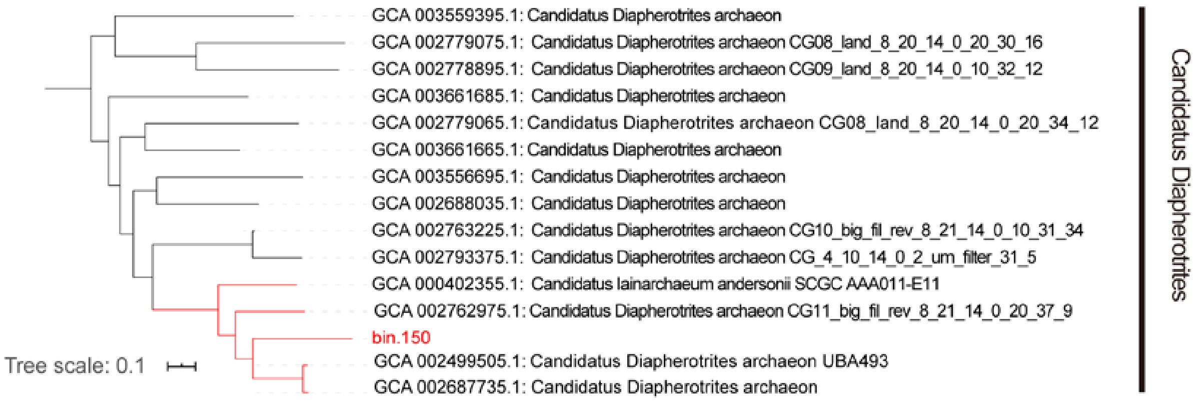


Figure S1. Phylogenetic analyses of bin.150 and some other Candidatus Diapherotrites archaeon reference genomes downloaded from ncbi database. A FastTree approximate maximum-likelihood phylogenetic tree was built using 15 conserved ribosomalprotein.

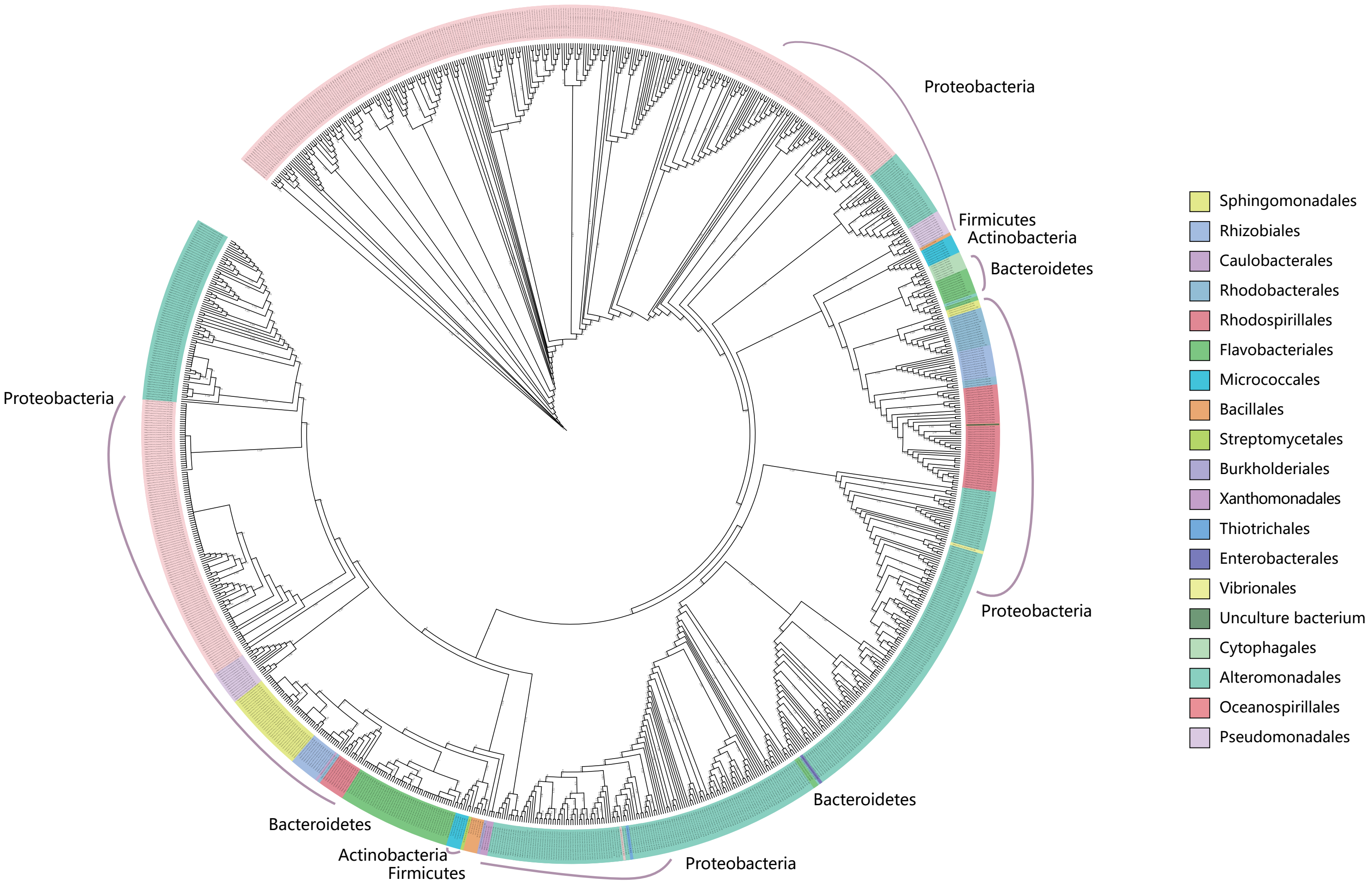


Figure S2. Phylogenetic analyses of 1070 bacterial isolates from the hadal sediment based on the 16S rRNA sequence with the FastTree. The color bar represents order level.

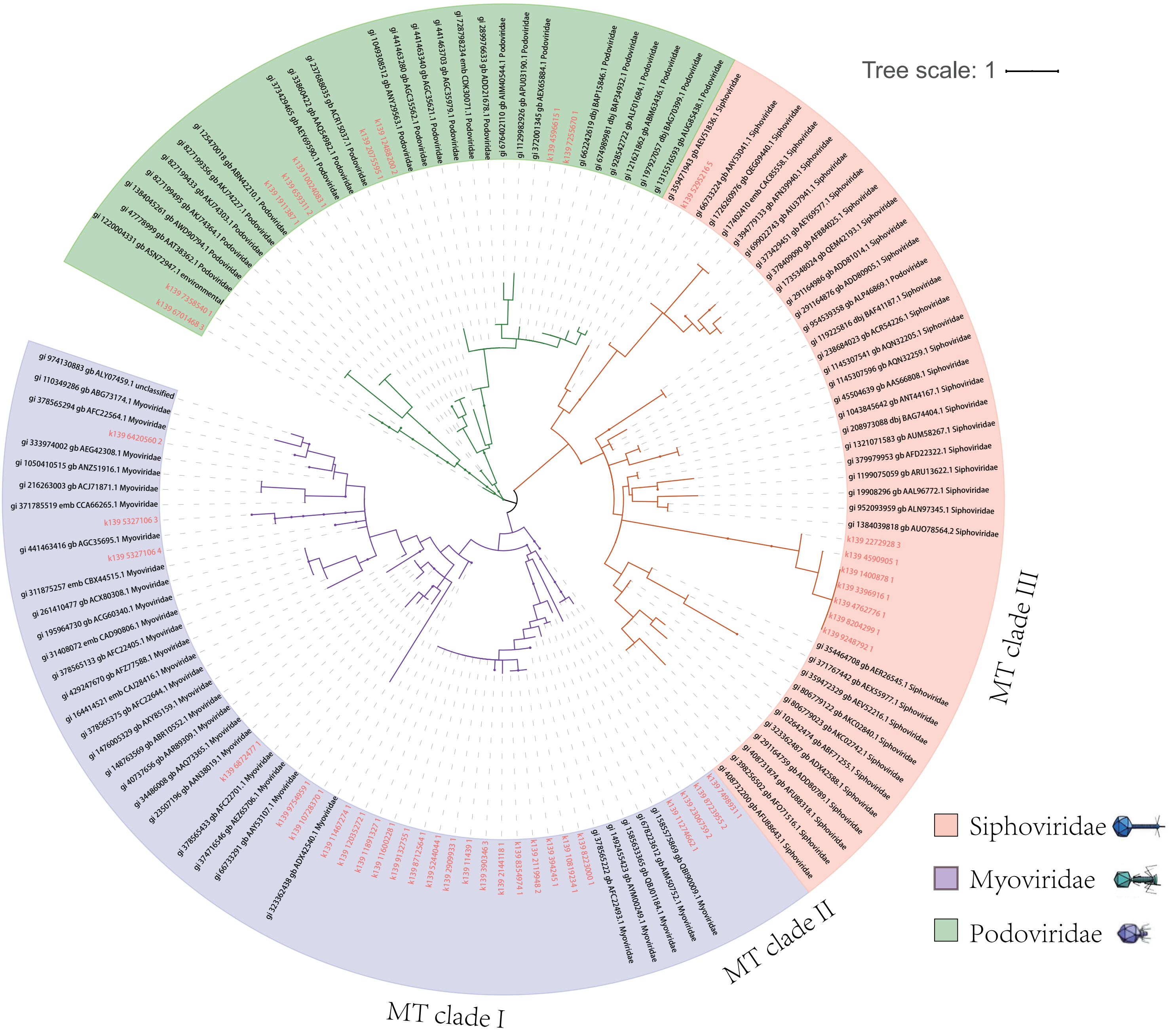


Figure S3. Phylogenetic analysis of Caudovirales based on TerL using the maximum likelihood algorithm. Reference viral sequences from NCBI are colored in black. Scale bar, one amino acid substitution per site.

A

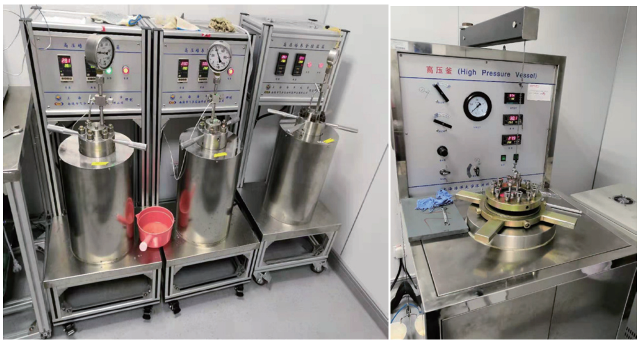


B

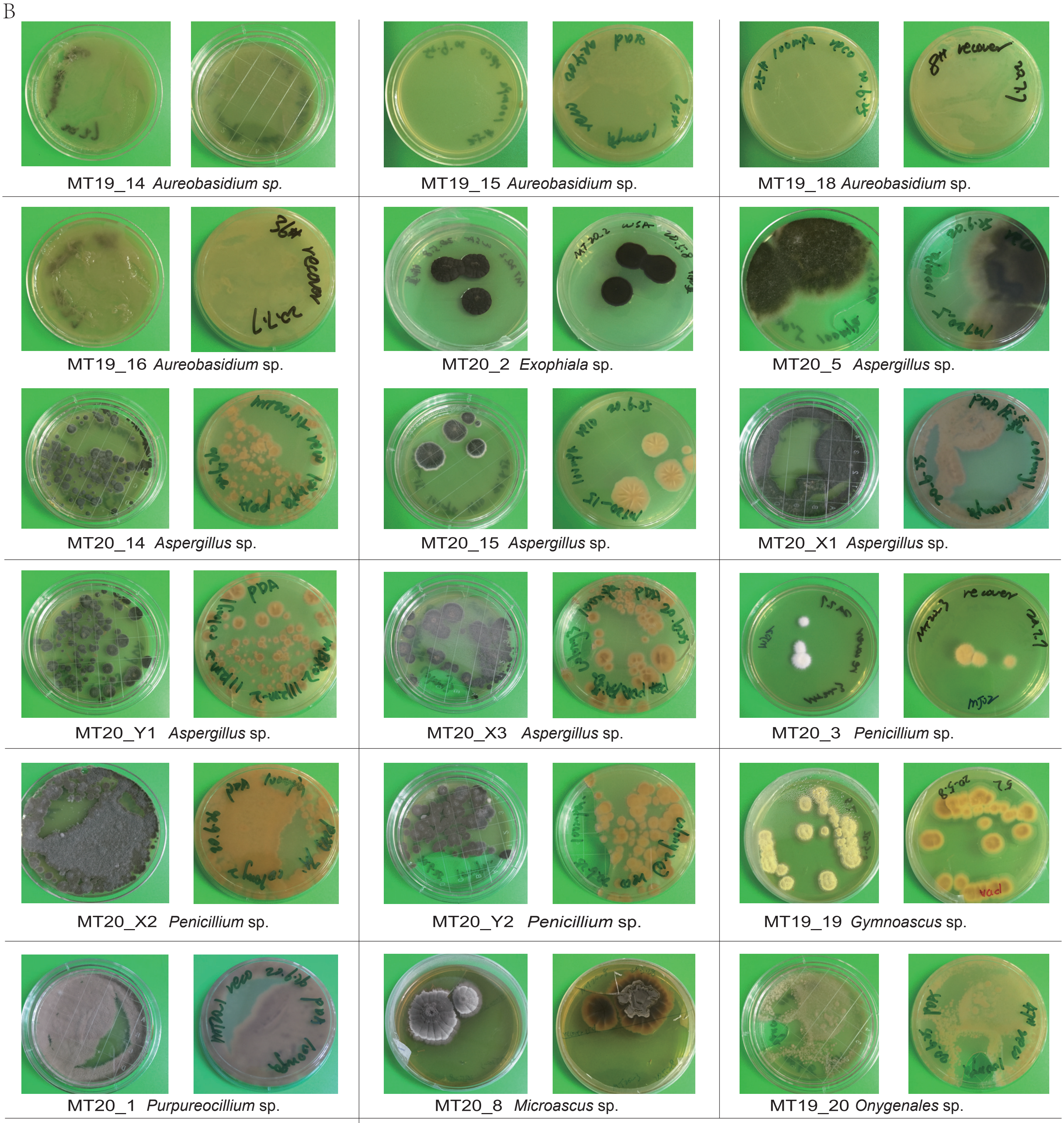


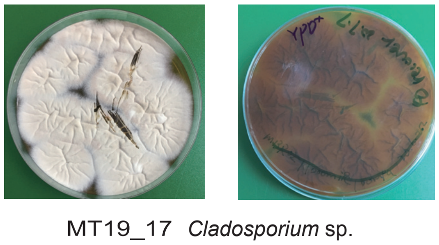


Figure S4. Culture of fungi under elevated hydrostatic pressure.
A) High pressure cultivation systerm.
B) Nineteen fungi isolated from the Challenger Deep sediment, which were tested to be viable to grow on plates after subjecting to high hydrostatic pressure (100 MPa) for fourteen days. Isolate names are listed on bottom.
