## Additional file 3 for "Revealing the full biosphere structure and versatile metabolic functions in the deepest ocean sediment of the Challenger Deep"

### Additional file 3: Analysis of MAGs on the potential for different types of fermentation

For microbes capable of anaerobic respiration, many were found to have the potential for different types of fermentation that degrade organic matter. The identification of elements of pyruvate reduction to fermentative end products in hadal MAGs, indicating the fermentative capabilities of hadal microbes. Key enzymes of pyruvate reduction, *PFOR/kor*, were distributed in 146 MAGs (24 phyla, Fig.2 and Additional file 1: Table S6). Marker genes for fermentation product metabolism, such as acetate, ethanol, and formate metabolism, were also found to be widely distributed in our hadal MAGs. The marker genes in acetate metabolism, i.e. (ADP-forming) acetate-CoA ligase (*Acd*), acetate kinase (*Ack*), and acetyl-CoA synthetase (*Acs*) were found in 144 MAGs (80%) associated with 22 phyla (Fig. 2 and Additional file 1: Table S6). *Acs* was the most widespread and also found in archaea like Thaumarchaeota and Ca. Woesearchaeota, indicating the role of archaea in fermentative acetate production in the hadal sediment. Ethanol metabolism, marked by aldehyde and alcohol dehydrogenases (*Aldh/Adh/Exa/Eut/Frm/Yia*) appeared to be the second most widespread in 16 phyla corresponding to 82 MAGs (46%). Formate metabolism, marked by formate dehydrogenase (*Fdh/Fdo*) was widely found in 67 MAGs (37%) associated with 13 phyla. These results indicated the importance of acetate, ethanol, and formate as fermentative metabolites for the hadal microbes.

Other fermentative metabolism-related genes, such as lactate dehydrogenase, phosphate acetyltransferase, and acylphosphatase, were also found scattering in the MAGs for the hadal sediment microbiome. Particularly, acylphosphatase (*acyP*) involved in anthranilate degradation, was widely distributed in 100 MAGs (56%; 24 phyla), indicating benzoate is an important fermentative metabolite in the hadal sediment, and the degradation of aromatic compounds is a common metabolic requirement for the Challenger Deep sediment microbes.
